# Pathway-Centric Integration of CRISPR Fitness with Molecular Features Draws Cancer State Maps

**DOI:** 10.64898/2026.05.12.724547

**Authors:** Tamaki Hagiwara, Kei K Ito, Kanako Kuroki, Ryuta Niikura, Masayasu Hirota, Kentaro Kawai, Hideko Uga, Junichi Iwakiri, Goro Terai, Kiyoshi Asai, Susumu Goyama, Shiori Shikata, Akinori Kanai, Yutaka Suzuki, Kenichi Shimada, Masamitsu Fukuyama, Shoji Hata, Shohei Yamamoto, Takumi Chinen, Daiju Kitagawa

## Abstract

Cancer cells display heterogeneous pathway activity that shapes therapeutic vulnerability, but mapping it remains challenging. Transcriptomic scores do not directly measure functional activity, and CRISPR knockout data alone lack molecular interpretability. We introduce StateMap, a pathway-centric framework integrating gene expression and genome-wide CRISPR knockout fitness data from the Cancer Dependency Map. For a given pathway, StateMap selects features by co-dependency and mutual information, then projects cell lines into a low-dimensional space reflecting pathway activity and molecular state. Applied to the Hippo pathway, it resolved five functional states refining the YAP-on/YAP-off dichotomy. Notably, the ‘Hippo-strong’ state showed selective dependence on integrin αVβ5; ITGAV depletion triggered Hippo-dependent cell aggregation and G1 arrest via enhanced cell–cell adhesion. Machine learning transfer to TCGA identified a matching subtype with poor prognosis, nominated NNMT as a biomarker, and predicted sensitivity to the αV inhibitor Cilengitide. StateMap enables pathway-specific state mapping and discovery of state-selective therapeutic vulnerabilities.

## Introduction

No two cancers are identical. Each harbors a unique combination of genetic, epigenetic, and transcriptional alterations that give rise to distinct patterns of pathway activity^1–3^. Because pathway activity strongly influences therapeutic response, precision oncology requires both the delineation of pathway state diversity across cancers and the identification of the molecular features that shape these states, enabling their interpretation and prediction in patients ^4^.

To date, pathway activity has been estimated primarily from transcriptomic data. Single-sample enrichment methods such as GSVA^5^ and ssGSEA^6^ compute per-sample scores from transcriptomic profiles and predefined signature gene sets, while footprint-based approaches such as PROGENy^7^ infer signaling activity from downstream transcriptional responses. Although these methods have proven valuable for stratifying tumors, they do not measure functional pathway activity directly ^8^. Functional activity is heavily modulated by post-translational modifications, protein stability, sub-cellular localization, and the availability of interacting partners ^9–11^. Therefore, whether a pathway component is highly transcribed and whether it is functionally active remains distinct questions.

The Cancer Dependency Map (DepMap) offers a complementary, function-oriented perspective. By profiling genome-wide CRISPR-knockout fitness effects across more than 1,000 cancer cell lines, DepMap directly measures the degree to which each gene is required for proliferation and survival ^12–14^. When a given oncogenic pathway is strongly active, knockout of its components markedly reduces fitness; the converse holds for tumor-suppressor pathways. Building on this principle, pathway-level representations can be derived from dependency data ^15^. For example, the Webster method decomposes the dependency matrix (cell lines × genes) into a dictionary (cell lines × latent functions) and a sparse loading matrix (latent functions × genes), where the dictionary values can be interpreted as the importance or activity of each latent function within each cell line ^16^. However, because such approaches operate on fitness data alone, the inferred pathway-level functional activity is not directly anchored to molecular features such as expression levels, thereby limiting biological interpretability.

Here, we introduce StateMap, a framework that maps functional pathway states across cancers by jointly leveraging gene expression and knockout fitness data (Fig. 1a). For a pathway of interest, StateMap selects input features through a data-driven approach. First, StateMap detects co-dependency modules from CRISPR knockout fitness profiles and selects genes from modules functionally associated with the pathway of interest. It then calculates mutual information (MI) to identify gene expression features most closely linked to these pathway-associated dependencies. StateMap subsequently integrates these features into a low-dimensional representation, projecting cell lines into a continuous space defined by both pathway activity and underlying molecular profiles. This approach enables direct visualization and comparative analysis of pathway states, which cannot be achieved using conventional scalar pathway scores alone.

**Figure 1.**
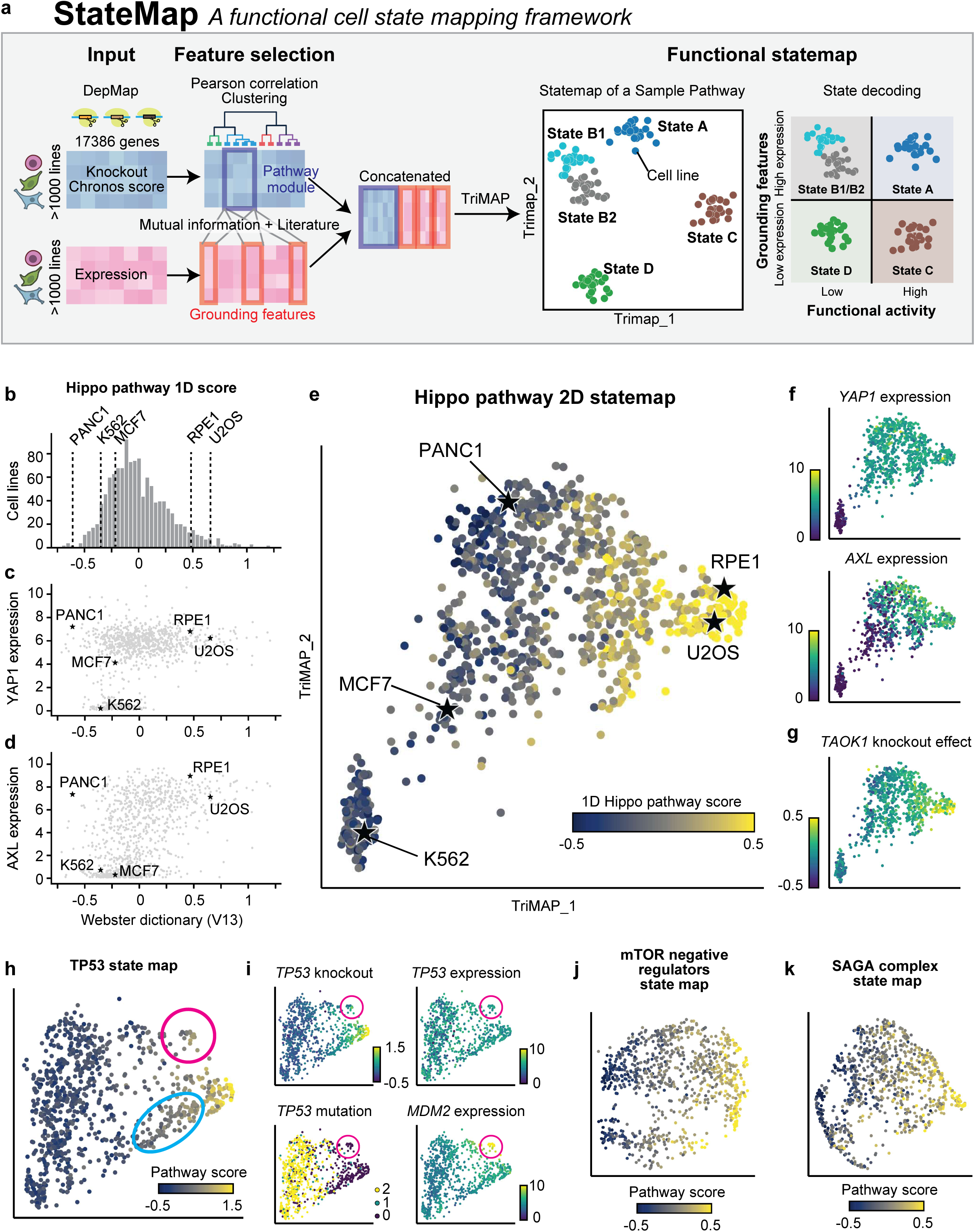
**(a)** Overview of the StateMap workflow. Genes composing pathway modules are detected based on knockout correlation clustering, and grounding features are selected based on mutual information and previous knowledge. The data are joined and normalized. Dimensionality reduction (TriMAP) is then performed for visualization, followed by state decoding to annotate each cell cluster. **(b)** Histograms showing the distribution of cell lines based on their Webster dictionary score (V13, calculated using DepMap 26Q1), corresponding to Hippo pathway activity. **(c)** Scatter plots showing the relationship between 1D Hippo activity scores and YAP1 expression levels (log_2_TPM). ‘RPE1’ denotes RPE1SS48 in DepMap, which exhibits a wild-type karyotype^65^. **(d)** Scatter plots showing the relationship between 1D Hippo activity scores and AXL expression levels (log_2_TPM). **(e)** Scatter plot showing the TriMAP embedding of cell lines based on Hippo-related gene features, with data points colored by the 1D pathway activity score from (b). **(f)** TriMAP embedding colored by expression levels of YAP1 and AXL (log_2_TPM). **(g)** TriMAP embedding colored by the knockout effect (Chronos score) of TAOK1. **(h)** Scatter plot showing the TriMAP embedding of cell lines based on TP53-related gene features, with data points colored by the corresponding 1D pathway activity score. **(i)** TriMAP embedding colored by the knockout effect (Chronos score) of TP53, expression levels of TP53 and MDM2 (log_2_TPM), and the mutation count of TP53. **(j)** Scatter plot showing the TriMAP embedding of cell lines based on negative mTOR regulator-related gene features, with data points colored by the corresponding 1D pathway activity score. **(k)** Scatter plot showing the TriMAP embedding of cell lines based on SAGA complex-related gene features, with data points colored by the corresponding 1D pathway activity score.

As a proof of concept, we applied StateMap to the Hippo signaling pathway, whose contribution to cell fitness varies widely across cancers. This analysis identified five distinct functional states corresponding to different modes of pathway status. Experimental validation in representative cell lines revealed that one specific state—characterized by hyperactive Hippo signaling—exhibits a unique vulnerability to integrin αV (ITGAV) inhibition. Subsequent integration with TCGA patient cohorts identified a matching tumor subtype associated with poor prognosis that may benefit from ITGAV-targeted therapy.

## Result

### Multi-dimensional integration by StateMap overcomes the limitations of single-axis pathway scores

To motivate the need for a multi-dimensional pathway representation, we first evaluated the limitations of one-dimensional pathway scores derived from DepMap dependency data via Webster^16^. As a representative case study, we focused on the Hippo signaling pathway, which displayed one of the most highly variable dependency patterns across cancer cell lines (11 of the 129 genes with the highest selectivity^17^—defined as >3 sigma from a normal distribution—belong to the Hippo pathway; Fig. S1a–e). In the Webster analysis, a specific latent factor (identified as column in the dictionary matrix) captured the Hippo pathway, with its values estimated based on the knockout effects of over 70 correlated genes, including *NF2*, *TAOK1*, and *LATS2* (Table S1, 2). The resulting Hippo pathway scores exhibited a unimodal, right-skewed distribution, indicating that a distinct subset of samples possessed notably elevated pathway activity (Fig. 1b). These cell lines (e.g., RPE1 and U2OS) demonstrated elevated expression of the pathway effectors YAP/TAZ and their downstream targets *AXL/CCN1/CCN2* (Fig. 1c, 1d, S1f-h)^18^, suggesting basally active YAP/TAZ. Conversely, the molecular underpinnings of the low-activity cell lines were considerably more diverse. For instance, K562 lacks YAP/TAZ expression altogether; MCF7 expresses YAP/TAZ but lacks downstream transcriptional activity; and PANC1 exhibits active YAP/TAZ (evidenced by the high expression of downstream targets) yet still displays low functional pathway dependency (Fig. 1c, 1d, S1f-h). When projected onto a one-dimensional axis, these molecularly distinct cell lines are distributed closely together, effectively obscuring their divergent molecular profiles (Fig. 1b). This compression illustrates that a single dimension is insufficient to capture such molecular complexity, thereby underscoring the necessity of integrating multiple features simultaneously to achieve accurate biological interpretation.

To address this limitation, we developed StateMap, a computational framework that integrates multi-dimensional knockout effects and molecular profiles—such as gene expression—into a low-dimensional latent space (Fig. 1a). Specifically, we selected knockout dependency profiles and expression levels of pathway-related genes and applied dimensionality reduction. Feature selection was rigorously performed to maximize both biological interpretability and functional relevance. For the dependency data, we selected genes exhibiting strong co-dependency, based on the rationale that highly correlated genes tend to share functional similarities ^14,15^ (Fig. S1a–e). This principle is also central to Webster, which leverages co-dependency-based regularization in its matrix factorization ^16^. For the transcriptomic data, we calculated the mutual information^19^ (MI) between gene expression and the knockout response to identify genes whose expression is strongly associated with this functional dependency (Fig. S2a). Unlike Pearson or Spearman correlation, which detect only linear or monotonic relationships, MI captures complex relationships, including non-linear and threshold-like behaviors. Notably, several previously reported Hippo downstream genes exhibited minimal MI values, suggesting that their effects are context-specific rather than universally applicable (Fig. S2a) ^20,21^.

Among the dimensionality reduction methods, TriMAP is specifically designed to capture non-linear relationships while preserving global structure ^22^, and we confirmed this advantage by measuring the global score (TriMAP^22^ 0.86, UMAP^23^ 0.55, t-SNE^24^ 0.78; Fig. S2b, c). This property was important for StateMap because biological interpretation depends on the faithful representation of relative distances among cell lines in the original dependency-expression feature space. Therefore, TriMAP was adopted for StateMap.

StateMap successfully resolved the heterogeneity of the Hippo pathway, distinctly segregating PANC1, MCF7, and K562 based on their integrated molecular-functional states while maintaining the spatial proximity of biologically similar cell lines, such as U2OS and RPE1 (Fig. 1e). The resulting embedding faithfully captured differences in Webster-derived Hippo pathway scores, transcriptomic levels, and knockout effects of related genes across the cell lines (Fig. 1e–g). To validate our data-driven gene selection strategy, we compared it against an embedding constructed from literature-based gene sets: 152 genes from the KEGG “Hippo signaling pathway” (hsa04390) for the Chronos input ^25^, and 22 genes from Wang et al.^20^ together with 80 genes from Kanai et al.^21^ for the expression input. This literature-based embedding produced a less distinct separation of cell lines along the Hippo pathway score axis, with cells from different functional states appearing intermixed (Fig. S2d, e). These results indicate that our combined co-dependency- and MI-based gene selection strategy resolves pathway states more effectively than curated gene lists alone.

To test StateMap’s generalizability, we applied the framework to several additional selective pathways, including *TP53*, the SAGA complex, and mTORC1 negative regulators (Fig. 1h–k, S1d, e, S2f). For the *TP53* pathway, we integrated the knockout effects of five core regulators (e.g., *TP53*, *ATM*) with the expression of four recognized targets (e.g., *CDKN1A*, *MDM2*) exhibiting high mutual information scores (Fig. S1e, S2f)^26^. The resulting embedding revealed clear variation in *TP53* pathway activity. As expected, many *TP53*-inactive cell lines harbored *TP53* mutations (Fig. 1h, i). Notably, however, a distinct subpopulation of *TP53* wild-type cell lines also exhibited low pathway activity (Fig. 1h). To investigate this discrepancy, we examined the expression levels of pathway-related genes. Among wild-type cell lines, *TP53* expression was comparable regardless of pathway activity, indicating that TP53 transcript levels do not explain the observed differences (Fig. 1i). Instead, the *TP53* wild-type, low-activity subpopulation (circled in magenta) displayed marked *MDM2* overexpression—a well-characterized mechanism of *TP53* pathway suppression (Fig. 1i)^27^. Other *TP53* wild-type but pathway-inactive subsets (circled in blue) may reflect additional, as-yet-uncharacterized inactivation mechanisms. Together, these findings demonstrate that StateMap provides a multi-dimensional representation of pathway status, anchored in specific molecular features, that single-score approaches fail to resolve. Below, we focus on the Hippo pathway to characterize the functional states resolved by this approach.

### Functional states of Hippo pathway clusters

To define interpretable Hippo pathway states, we applied spectral clustering to the StateMap affinity graph^28^. Analysis of the normalized graph Laplacian revealed a pronounced eigengap between the fifth and sixth smallest eigenvalues, indicating five dominant partitions (Fig. S3a, b)^29^. Importantly, this five-state structure was also recovered in the original dependency-expression feature space, confirming that it reflects intrinsic biological organization rather than an artifact of dimensionality reduction (Fig. S3c, d). The robustness of K = 5 was further validated by repeated subsampling, which produced a block-diagonal consensus matrix and a locally minimal PAC score (0.021)—substantially lower than those obtained at K = 4 (0.037) or K = 6 (0.073) (Fig. S3e, f)^30,31^. We therefore adopted these five clusters as the basis for downstream biological interpretation (Fig. 2a, b, Table S3).

**Figure 2.**
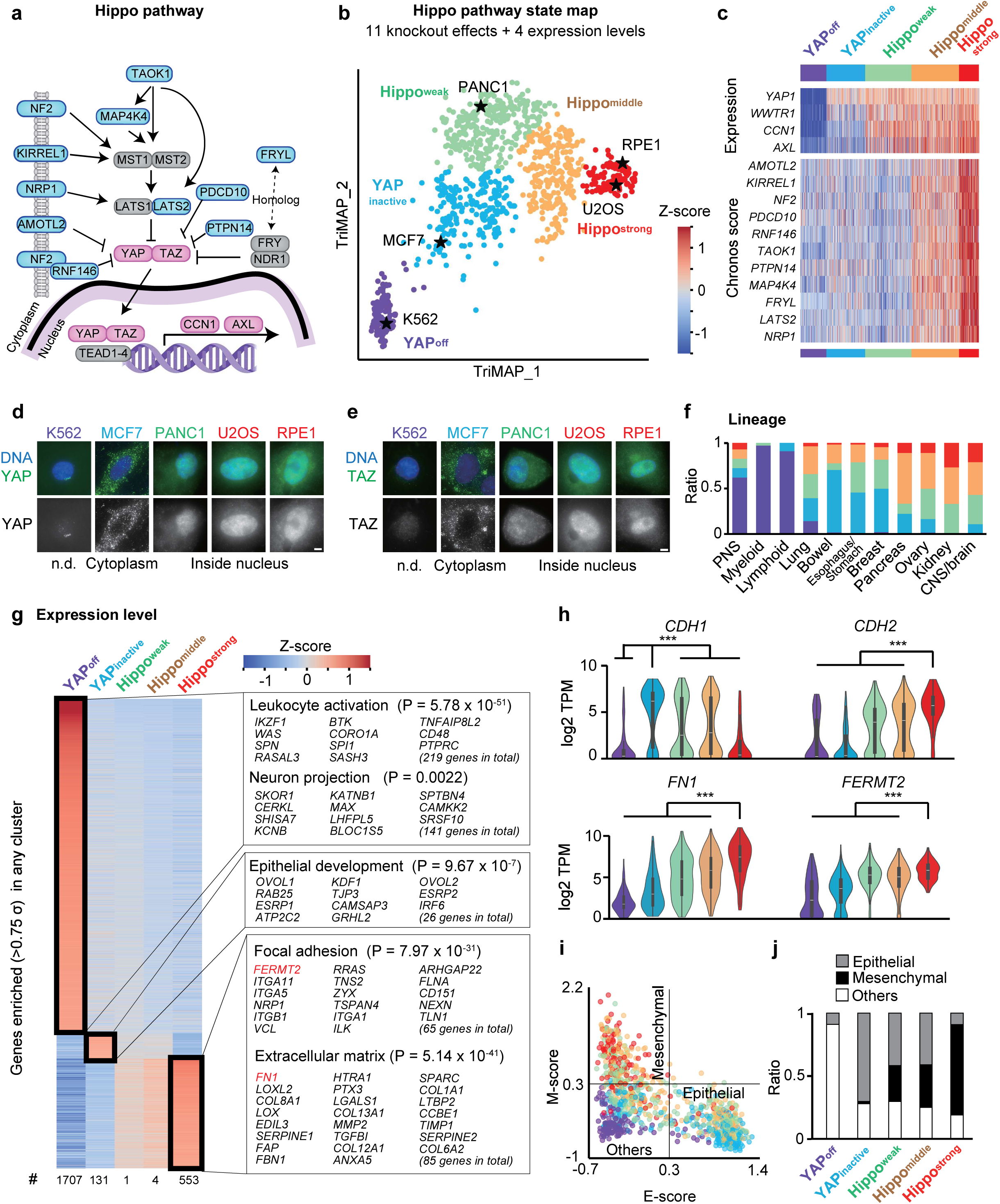
**(a)** Schematic representation of the Hippo signaling pathway. Knockout effect data for 11 genes (blue) and expression data for 4 genes (pink) are used in the dimensionality reduction. **(b)** Scatter plot showing the TriMAP embedding of cell lines based on Hippo-related gene features, with data points colored by cluster identity. **(c)** Heatmap showing expression levels and Chronos scores used for the dimensionality reduction. **(d)** Immunofluorescence images of YAP signals for the cell lines in (b). **(e)** Immunofluorescence images of TAZ signals for the cell lines in (b). **(f)** Distribution of Hippo-derived clusters across different cell types. **(g)** Gene expression profiles for genes enriched by >0.75σ and FDR < 0.01 in any of the five clusters. Genes within the rectangle are annotated with the corresponding enriched GO terms and their P values. **(h)** Violin plots showing the distribution of expression levels across the five clusters for representative genes. *P* values were calculated using the two-tailed Mann-Whitney U test. \*\*\**P*□<□0.001. **(i)** Scatter plot showing the E-score and M-score of each cell line. **(j)** Distribution of epithelial, mesenchymal, and other cancer types across different clusters.

The five distinct groups are defined by a unique combination of gene knockout effects and expression profiles. Two clusters correspond to states of weak YAP/TAZ activity, characterized by low target gene expressions including *CCN1/AXL* (Fig. 2c, S4a)^20^. Specifically, the ‘YAP-off’ cluster (e.g., K562) is defined by minimal YAP/TAZ expression, whereas the ‘YAP-inactive’ cluster (e.g., MCF7) expresses YAP/TAZ but sequesters the proteins in the cytoplasm, preventing target gene expression (Fig. 2c-e, S4a). The remaining three clusters exhibit active YAP/TAZ—marked by nuclear localization and high target gene expression—but differ in their sensitivity to Hippo pathway perturbation (Figs. 2c-e, S4a). This sensitivity gradient ranges from ‘Hippo-weak’ cells (e.g., PANC1), which are insensitive to Hippo gene perturbation, to ‘Hippo-strong’ cells (e.g., U2OS and RPE1), which gain a profound growth advantage from Hippo pathway gene knockout, with a ‘Hippo-middle’ group showing intermediate sensitivity (Fig. 2c). Together, the five clusters provide a functional atlas of Hippo pathway states across cancer cell lines.

Analysis of cancer lineages and subtypes revealed that Hippo pathway state often transcends traditional classification boundaries. While certain patterns were observed—for example, peripheral nervous system (PNS), myeloid, and lymphoid cancers were frequently YAP-off, and bowel-derived cancers tended to be YAP-inactive—most lineages displayed considerable heterogeneity (Fig. 2f, S5a, b). At the subtype level, neuroendocrine small cell lung cancer consistently exhibited a YAP-off state, and uterus endometrial carcinoma showed a Hippo-weak signature (Fig. S6). Nevertheless, the majority of cancer subtypes remained highly heterogeneous with respect to Hippo pathway status (Fig. S6).

Notably, prior studies defined cancers as YAP-on or YAP-off based on YAP/TAZ expression, identifying most solid tumors as YAP-on and most hematological, neural, or neuroendocrine cancers as YAP-off ^32,33^. Our five-cluster model refines this binary framework by subdividing the YAP-on population into four functional groups, providing a more detailed atlas of Hippo pathway states.

### Transcriptional characterization of Hippo clusters

To further characterize these clusters over the frame of cell type, we analyzed genes uniquely upregulated in each cluster by comparing median expression values between one cluster and all others. The ‘YAP-off’ cluster showed the highest number of uniquely upregulated genes, many of which, as expected from its lineage composition, were related to leukocyte, neural, and neuroendocrine functions ^34,35^ (Fig. 2g, S5c). The remaining four clusters exist along an epithelial-to-mesenchymal spectrum. At one end, the ‘YAP-inactive’ cluster is distinctly epithelial, enriched for epithelial markers such as E-cadherin (*CDH1*) among its uniquely upregulated genes (Fig. 2g, h). At the other end, ‘Hippo-strong’ cells are highly mesenchymal, significantly upregulating mesenchymal markers such as N-cadherin (*CDH2*) and extracellular matrix components (e.g., *FN1, FBN1, COL8A1*), along with focal adhesion/integrin-signaling genes (e.g., *FERMT2, ILK*) (Fig. 2g, h). The ‘Hippo-weak’ and ‘Hippo-middle’ clusters occupy an intermediate state with few uniquely upregulated genes. Consistently, pan-cancer EMT signature scoring classified ‘YAP-inactive’ cells as predominantly epithelial and ‘Hippo-strong’ cells as mesenchymal, whereas ‘Hippo-weak’ and ‘Hippo-middle’ cells showed heterogeneous intermediate profiles^36,37^ (Fig. 2i, j). These results indicate that EMT status represents an important transcriptional axis within the Hippo StateMap, but that the five-state structure cannot be reduced to EMT alone. Given their distinctive expression profiles and strong proliferative response to Hippo-gene knockout, subsequent analyses focused on the Hippo-strong cells.

### Activated cell-matrix adhesion via integrin αVβ5 is the hallmark of the Hippo-strong state

To identify the molecular basis of the Hippo-strong phenotype, the knockout effect profiles were analyzed. By definition, 11 selective Hippo pathway genes exhibited relatively high Chronos scores in Hippo-strong cells, accompanied by other Hippo-related genes (e.g., *SAV1, TAOK2, RAP2B*) (Fig. 3a). In contrast, genes exhibiting strong dependency were enriched for focal adhesion components, including intracellular scaffold proteins (e.g., *FERMT2, PXN*) and focal adhesion–associated kinases (e.g., focal adhesion kinase (*PTK2*), integrin-linked kinase (*ILK*)) (Fig. 3a, b). Within focal adhesion complexes, integrins serve as transmembrane receptors that physically link extracellular ligands to the intracellular adhesion machinery^38^. Among integrins, only *ITGAV* (integrin αV) and *ITGB5* (integrin β5) emerged as selectively essential (Fig. 3c, d). These subunits form the αVβ5 heterodimer, which recognizes RGD-containing extracellular ligands^39^. Although αVβ5 integrin also participates in a noncanonical process termed reticular adhesion or flat clathrin lattice ^40,41^, other components of that pathway (e.g., clathrin, clathrin adaptors, stonin1 ^41^) were not specifically required in Hippo-strong cells (Fig. S5d). Moreover, fibronectin, which is highly expressed in Hippo-strong cells (Fig. 2), has been reported to suppress noncanonical adhesion formation ^42^. These findings suggest that αVβ5-containing canonical focal adhesions is crucial for cell growth of Hippo-strong cells.

**Figure 3.**
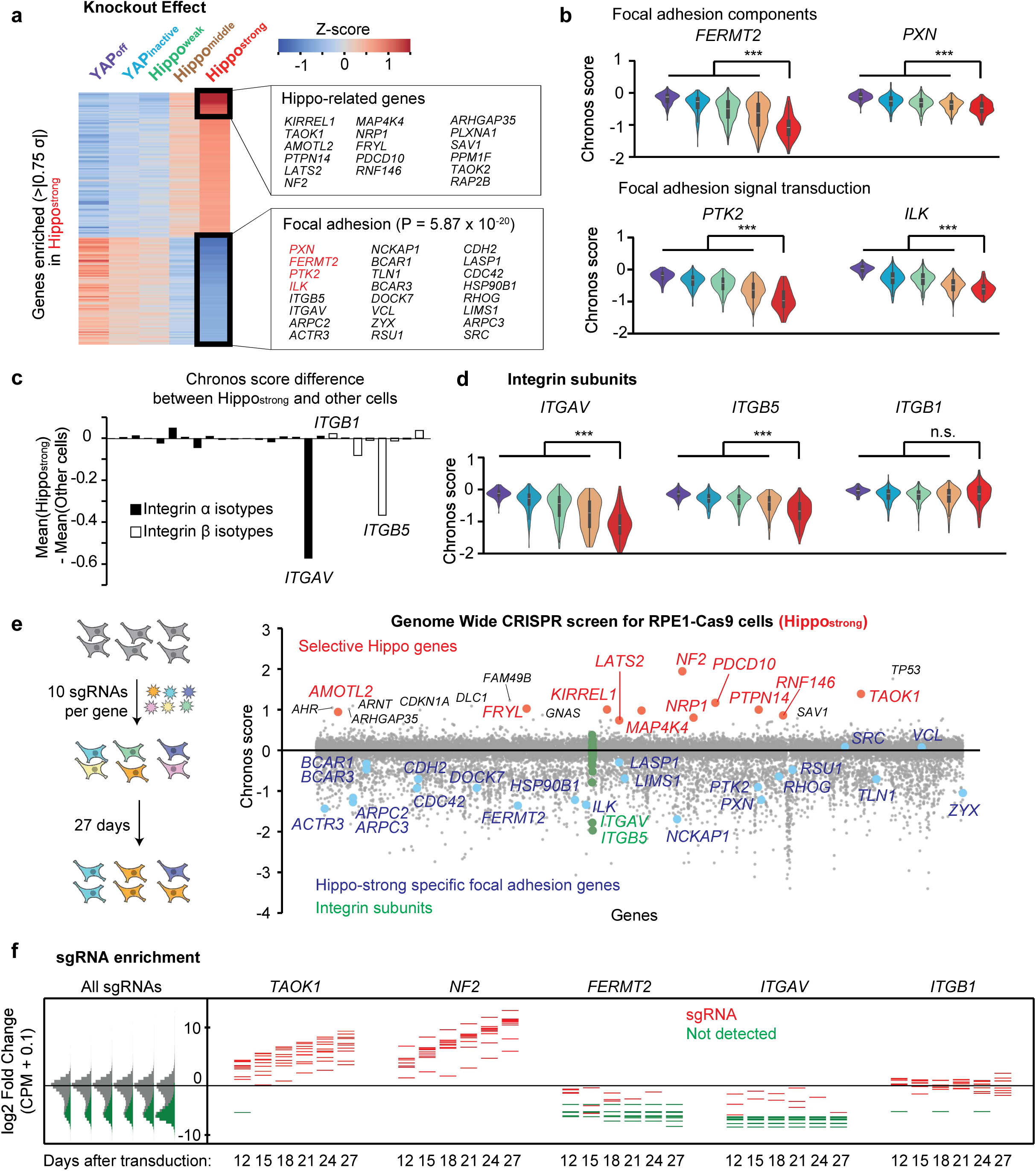
**(a)** Gene knockout effect profiles for genes enriched by >|0.75 σ| and FDR < 0.01 in Hippo-strong cells. Genes within the rectangle are annotated with the corresponding enriched GO terms and their P values. **(b)** Violin plots showing the distribution of expression levels across the five clusters for representative genes with low Chronos scores (i.e., higher essentiality) in the Hippo-strong cluster. *P* values were calculated using the two-tailed Mann-Whitney U test. \*\*\**P*□<□0.001. **(c)** Bar plot showing differences in Chronos scores of integrin coding genes between the Hippo-strong cluster and other cells. **(d)** Violin plots showing the distribution of Chronos scores of representative integrin genes across the five clusters. *P* values were calculated using the two-tailed Mann-Whitney U test. \*\*\**P*□<□0.001. n.s. P > 0.05. **(e)** Scatter plot showing CRISPR screen results, with selective Hippo-related genes highlighted in red, Hippo-strong–specific focal adhesion genes in blue, and integrin subunits in green. Genes are sorted alphabetically. **(i)** Log₂ fold changes of (counts per million + 0.1) across 10 sgRNAs for the indicated genes from day 12 to day 27. Undetected sgRNAs are shown in green.

To independently validate the knockout effects of Hippo-pathway and focal adhesion genes, we performed a genome-wide CRISPR–Cas9 screen in RPE1 cells, a representative ‘Hippo-strong’ model, using a high-coverage library (10 sgRNAs per gene ^43^). Chronos analysis confirmed that all 11 selective Hippo genes ranked within the top 0.15% of positively selected hits (growth advantage upon knockout; Fig. 3e, f; Tables S4–S5), whereas most of the focal-adhesion genes on which Hippo-strong cells depend (e.g., *FERMT2, PXN, PTK2, ILK*) were negatively selected (Fig. 3e, f). Among integrins, disruption of αV or β5 showed markedly stronger growth suppression than any other integrin subunits (Fig. 3e). Collectively, these observations—upregulation of ECM and focal adhesion genes, dependence on focal adhesion components, and specific reliance on αVβ5—identify hyperactive αVβ5-integrin–mediated focal adhesion signaling as a defining feature of the Hippo-strong state.

### ITGAV depletion triggers Hippo-dependent aggregation and G1 arrest in Hippo-Strong Cells

Comparative analysis suggested that integrin αVβ5–mediated focal adhesion is selectively important in ‘Hippo-strong’ cells. We therefore knocked down integrin αV (*ITGAV*) using siRNA in representative cells from different clusters and compared their phenotypes. In Hippo-strong cells (RPE1 and U2OS), *ITGAV* depletion caused cells to cluster together, forming compact aggregates (Fig. 4a-e). By contrast, this aggregation response was not observed in other cell types, including DLD1 and MCF7 from the YAP-inactive cluster or PANC1 from the Hippo-weak cluster (Fig. 4c–e), although they expressed comparable *ITGAV* and *ITGB5* (Fig. S7a).

**Figure 4.**
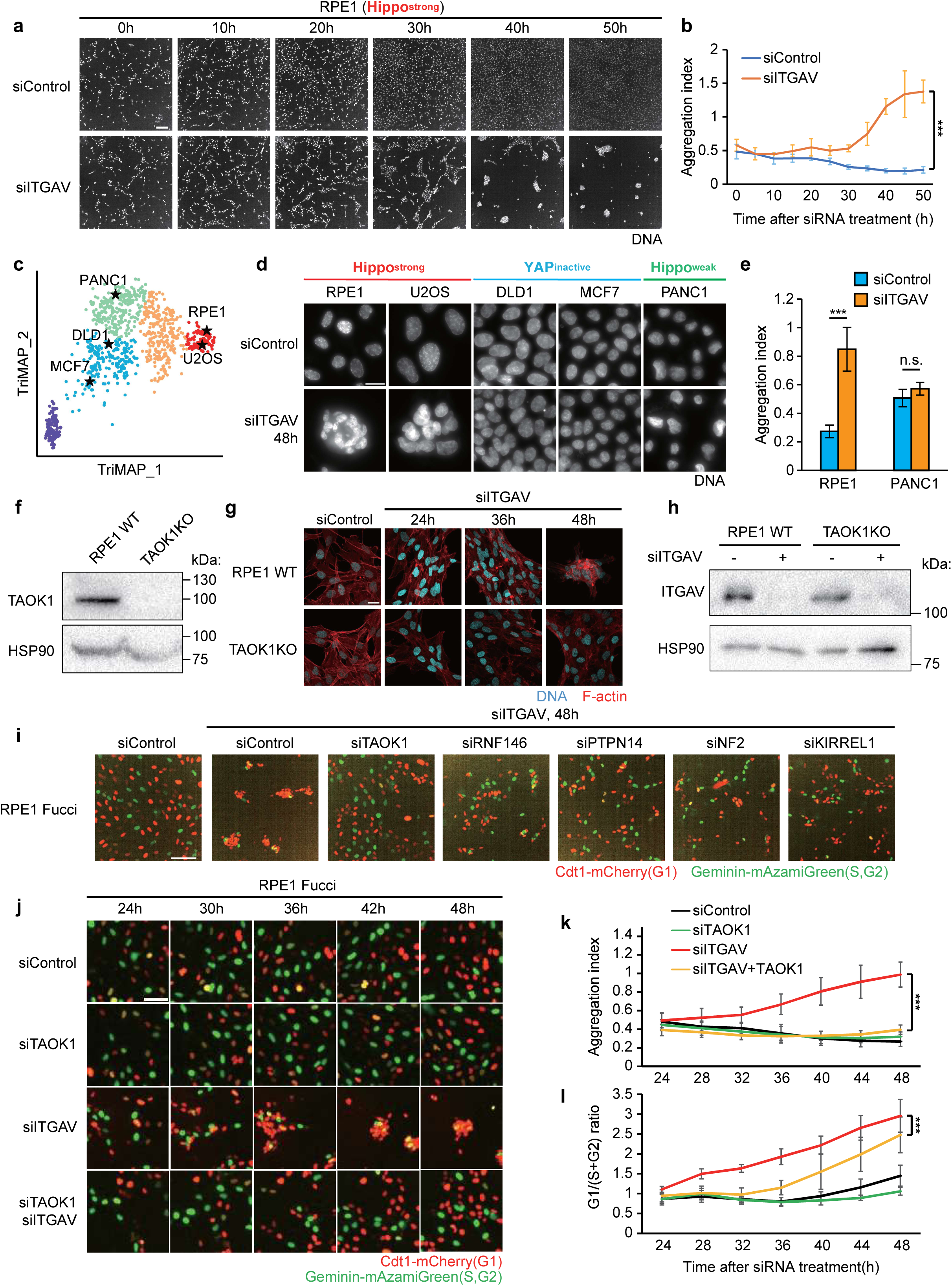
**(a)** Time-lapse images from live-cell imaging of RPE1 cells (Hippo-strong) treated with siControl or siITGAV. Scale bar, 100μm. **(b)** Quantification of cell aggregation in (a) over a time course from 5 to 55 hours post-siRNA transfection. *P* value was calculated for the area under the curves using the two-tailed Student’s *t-*test. \*\*\**P*□<□0.001. **(c)** TriMAP embedding highlighting the cell lines used in panel (d): RPE1 and U2OS (Hippo-strong), DLD1 and MCF7 (YAP-inactive), and PANC1 (Hippo-weak). **(d)** Immunofluorescence images of the cell lines in (c) treated with siControl or siITGAV. Scale bar, 100μm. **(e)** Quantification of cell aggregation in (d). *P* value was calculated for the area under the curves using the two-tailed Student’s *t-*test with Holm’s adjustment. \*\*\**P*□<□0.001; n.s., P > 0.05. **(f)** Western blot showing knockout efficiency of TAOK1 in RPE1 cells. **(g)** Time-lapse images of RPE1 wild-type (WT) and TAOK1 knockout (KO) cells treated with siControl or siITGAV. Scale bar, 100μm. **(h)** Western blot showing depletion efficiency of ITGAV in the cells used in (g). **(i)** Live-cell fluorescence microscopy images of RPE1-Fucci cells at 48 hours post-transfection with siITGAV and siRNAs targeting selective Hippo genes. Scale bar, 100μm. **(j)** Time-lapse images of RPE1-Fucci cells treated with the indicated siRNAs. Scale bar, 100μm. **(k)** Quantification of cell aggregation in (i) from 24 to 48 hours post-siRNA transfection. *P* value was calculated for the area under the curves using the two-tailed Student’s *t*-test. \*\*\**P*□<□0.001. **(l)** Quantification of the G1/(S+G2) cell ratio in RPE1-Fucci cells. *P* value was calculated for the area under the curves using the two-tailed Student’s *t*-test. \*\*\**P*□<□0.001.

We hypothesized that the aggregation phenotype requires a Hippo pathway activity characteristic of ‘Hippo-strong’ cells. To test this, *TAOK1*, a key Hippo pathway component, was deleted in RPE1 cells using a CRISPR-del strategy that removes the entire gene locus ^44^ (Fig. 4f, S7b-d). Strikingly, *ITGAV* depletion in these *TAOK1*-knockout cells failed to induce cell aggregation (Fig. 4g, h). Supporting this observation, siRNA-mediated knockdown of other selective Hippo pathway components including *RNF146*, *PTPN14*, *NF2*, and *KIRREL1* also effectively suppressed the aggregation phenotype (Fig. 4i). Together, these results indicate that the cell aggregation observed upon *ITGAV* depletion requires Hippo pathway activity.

To further understand the consequences of cell aggregation, we investigated the fate of the aggregated cells. While *ITGAV* depletion did not induce apoptosis (Fig. S7e), time-lapse microscopy with the Fucci cell cycle reporter system revealed that the aggregated cells arrested in the G1 phase, as indicated by the accumulation of the G1 marker Cdt1-mCherry (Fig. 4j-l). Suppression of aggregation through simultaneous knockdown of Hippo pathway genes significantly delayed the onset of G1 arrest (Fig. 4j–l), indicating that cell aggregation contributes to the cell cycle arrest. Conversely, directly inducing G1 arrest with a CDK4/6 inhibitor did not cause cell aggregation (Fig. S7f), confirming that the G1 arrest is not a cause of the aggregation. Taken together, these findings demonstrate that in Hippo-strong cells, the disruption of ITGAV-mediated adhesion triggers a Hippo pathway-dependent response characterized by cell aggregation that subsequently leads to G1 cell cycle arrest.

### Hippo pathway enhances cell-cell adhesion independent of YAP/TAZ

We next examined why Hippo pathway–suppressed cells, such as *TAOK1*-knockout cells, failed to undergo cell aggregation upon *ITGAV* depletion. Given that the canonical Hippo pathway suppresses YAP/TAZ activity, it seemed plausible that activated YAP/TAZ in *TAOK1*-knockout cells might prevent aggregation. However, even when YAP/TAZ was inhibited by siRNA or a pharmacological inhibitor, *TAOK1*-knockout cells still failed to aggregate (Fig. 5a–c). These results indicate that the Hippo pathway regulates cell aggregation independently of canonical YAP/TAZ regulation.

**Figure 5.**
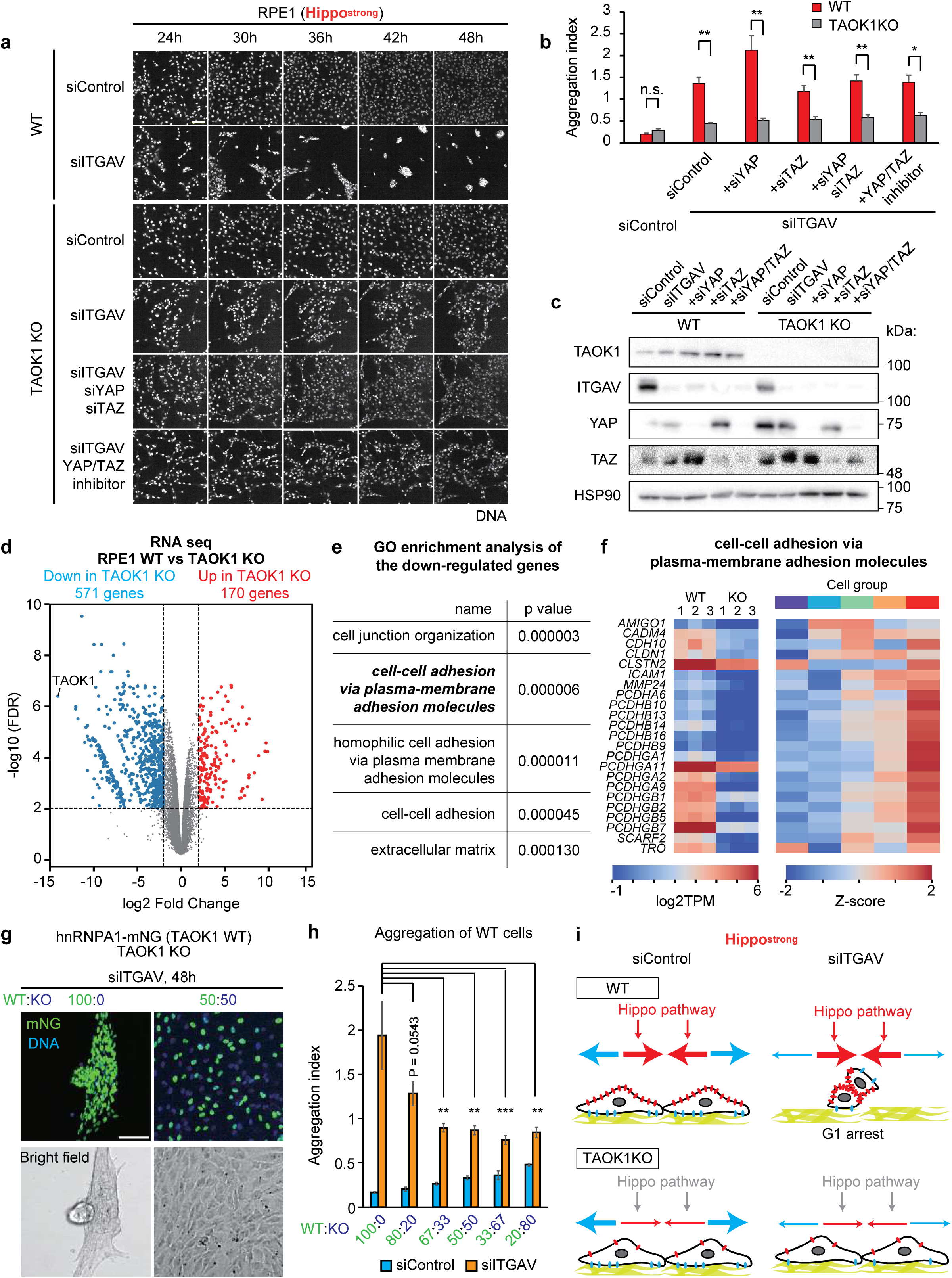
**(a)** Time-lapse images of RPE1 wild-type (WT) and TAOK1 knockout (KO) cells treated with the indicated siRNAs or compound. Scale bar, 100μm. **(b)** Quantification of cell aggregation in (a) at 48 hours post-siRNA transfection. *P* values were calculated using the two-tailed Student’s *t*-test with Holm’s adjustment. \*\**P < 0.01; *P < 0.05; n.s., P > 0.05*. **(c)** Western blot showing the depletion efficiency of ITGAV, YAP, and TAZ in the cells used in (a). **(d)** Volcano plot showing differential gene expression between WT and TAOK1 KO RPE1 cells. Upregulated genes are shown in red; downregulated genes in blue. **(e)** Gene Ontology (GO) enrichment analysis of the genes downregulated in TAOK1 KO cells compared to WT cells. **(f)** Heatmaps showing expression levels of downregulated genes associated with the GO term “cell-cell adhesion via plasma membrane adhesion molecules” in triplicate WT and KO samples (left), and across the five clusters defined in Figure 2 (right). **(g)** Immunofluorescence images of mixed RPE1 cells: hnRNPA1–mNeonGreen–expressing cells (TAOK1 wild type) and TAOK1 knockout cells, treated with siITGAV. Scale bar, 100μm. **(h)** Quantification of cell aggregation in (g). *P* values were calculated using the two-tailed Dunnett’s test. \*\*\**P < 0.001; **P < 0.01; n.s., P > 0.05*. **(i)** Schematic model of the observed phenotype. In Hippo-strong cells, both cell-cell adhesion and cell-matrix adhesion are elevated, and ITGAV depletion induces cell aggregation. In TAOK1 KO cells, reduced cell-cell adhesion prevents aggregation following ITGAV depletion.

To explore an alternative mechanism, we performed comparative RNA sequencing on wild-type and *TAOK1*-knockout RPE1 cells (Fig. 5d). Differential expression analysis identified 170 upregulated and 571 downregulated genes in knockout cells (FDR < 0.01, log2 fold change > 2 or < –2), with both groups enriched for cell-surface molecules (Fig. 5d, e, S8a, b; Table S6, S7). Notably, cell–cell adhesion molecules such as protocadherins were significantly overrepresented among downregulated genes (Fig. 5e, f), suggesting that impaired adhesion may underlie the reduced aggregation phenotype of knockout cells. To test this possibility, we conducted co-culture experiments with wild-type and knockout cells. If decreased aggregation in knockout cells results from reduced cell–cell adhesion, these less adhesive knockout cells would act as insulators and limit the aggregation of wild-type cells. Indeed, mixing in knockout cells limited the aggregation of wild-type cells, and this effect became more pronounced as the proportion of knockout cells increased (Fig. 5g, h). These results suggest that weakened cell–cell adhesion accounts for the reduced aggregation phenotype of *TAOK1*-knockout RPE1 cells. We therefore propose that the Hippo pathway, including *TAOK1*, normally promotes the expression of cell–cell adhesion genes in wild-type RPE1 cells, making them prone to aggregation.

Importantly, the adhesion molecules downregulated in *TAOK1*-knockout cells were expressed at higher levels in Hippo-strong cells compared with other clusters (Fig. 5f), suggesting that this pro-adhesion role extends beyond RPE1 cells. Together, these findings support a Hippo-strong model in which the active Hippo pathway enhances cell–cell adhesion but is counterbalanced by strong integrin αVβ5-mediated focal adhesion to maintain normal morphology (Fig. 5i). When this balance is disrupted by weakening focal adhesion, Hippo-induced adhesion predominates, leading to cell aggregation and G1 arrest (Fig. 5i).

### Machine learning identifies a Hippo-strong tumor subtype with a targeted therapeutic vulnerability

To assess the clinical relevance of the Hippo pathway states, we projected these states onto tumors from TCGA, TARGET, and Treehouse using a machine learning classifier^45^. To mitigate confounding effects from stromal and immune cells present in tumors but absent in cell lines, we used Celligner-aligned DepMap and tumor expression data (Fig. 6a)^45,46^. The aligned DepMap data was employed to train the machine learning classifier to predict Hippo pathway states (Fig. 6a). The classifier achieved a balanced accuracy of approximately 0.7, significantly outperforming a baseline random classifier (score of 0.2) (Fig. S9a, b). When applied to the aligned tumor expression data ^45^, the predicted Hippo states displayed tissue-specific tendencies similar to those observed in DepMap cell lines (Fig. 2f, 6b, S9c, Table S8). Moreover, consistent with our cell line findings (Fig. 5f), tumors classified as Hippo-strong exhibited higher expression of cell–cell adhesion molecules (Fig. 6c).

**Figure 6.**
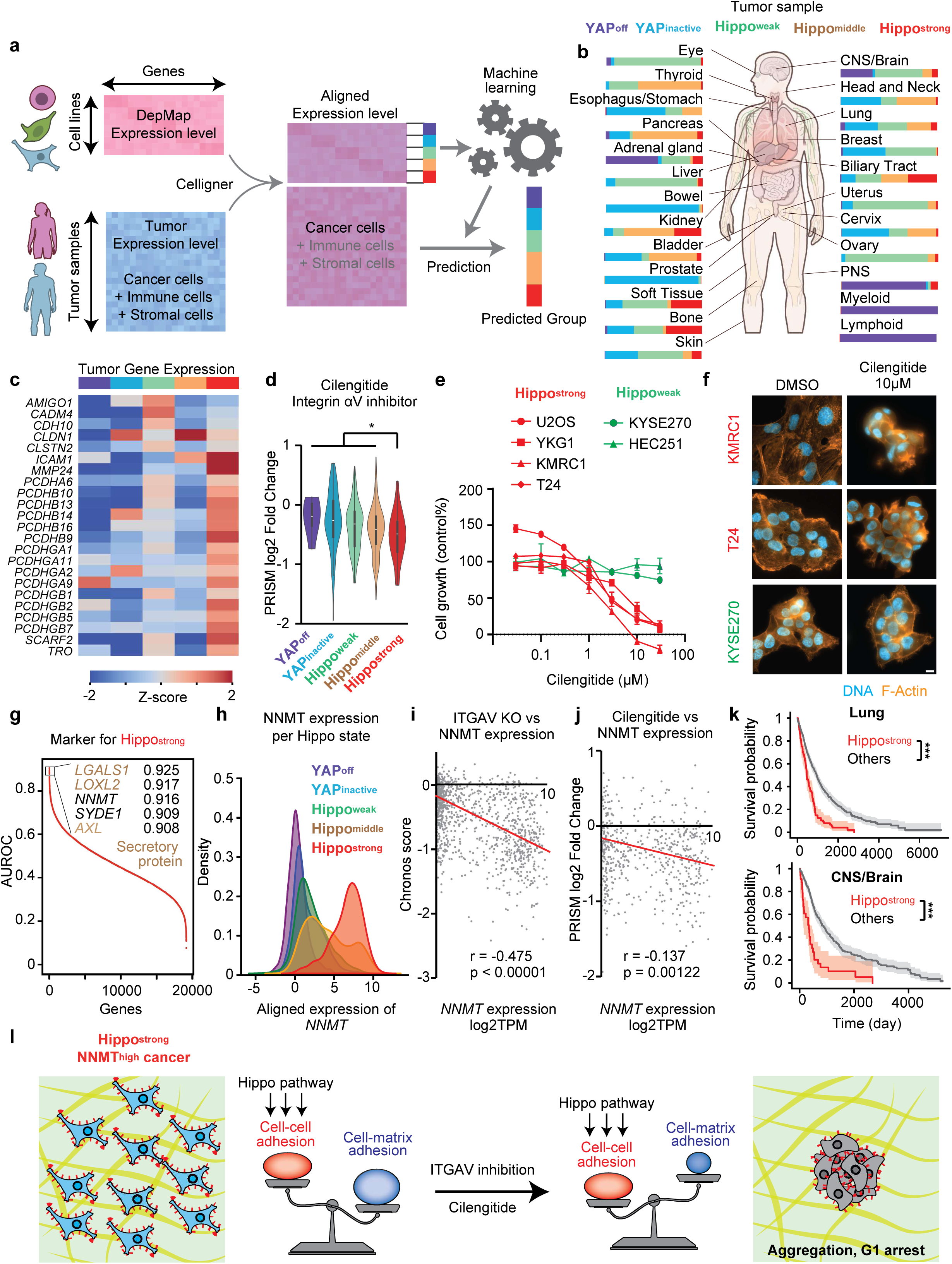
**(a)** Overview of the machine learning–based tumor classification workflow. Gene expression profiles of DepMap cell lines and tumor samples are aligned using Celligner to mitigate confounding effects from stromal and immune cell contributions in tumor samples. The aligned DepMap expression data is used to train a machine learning model to predict Hippo-derived clusters, which is then applied to classify tumor samples. **(b)** Distribution of predicted Hippo-derived clusters across different cancer types. **(c)** Heatmap showing expression levels of cell-adhesion related genes across the five clusters. **(d)** Violin plots showing the distribution of PRISM drug sensitivity scores for Cilengitide across the five clusters. *P* value was calculated using the two-tailed Mann-Whitney U test. \**P*□<□0.05. **(e)** Line plot showing the dose–response relationship between Cilengitide concentration and cell growth. **(f)** Immunofluorescence images of cells treated with 10 µM Cilengitide. Scale bar, 10μm. **(g)** Scatter plot showing the area under the ROC curve (AUROC) for each gene’s ability to distinguish Hippo-strong tumors based on aligned expression data. **(h)** Kernel density plot of *NNMT* expression levels in the TCGA dataset, colored by Hippo state cluster. **(i)** Scatter plot showing the relationship between *NNMT* expression levels and *ITGAV* Chronos score. P value was calculated using a two-tailed t-test on the regression slope. **(j)** Scatter plot showing the relationship between *NNMT* expression levels and PRISM log2 fold change of Cilengitide. P value was calculated using a two-tailed t-test on the regression slope. **(k)** Kaplan–Meier survival curves for lung cancers, comparing patients classified into the Hippo-strong cluster versus other clusters. *P* values were calculated using the log-rank test with Benjamini–Hochberg correction. \*\*\**P*□<□0.001. **(l)** Schematic model of Hippo-strong tumors. These tumors are characterized by high *NNMT* expression, increased cell–cell adhesion, and enhanced sensitivity to integrin αV inhibition.

To identify therapeutic vulnerabilities in this subtype, we analyzed drug sensitivity data from the PRISM screen^47^. As expected from the selective dependency on *ITGAV* in Hippo-strong cells (Fig. 3, 4), the integrin αVβ5/αVβ3 inhibitor Cilengitide ^48,49^ (Cyclic RGD peptides) was selectively effective compared to other clusters (Fig. 6d). We validated this finding by treating several Hippo-strong and Hippo-weak cell lines with Cilengitide, which selectively suppressed the growth of the Hippo-strong cells (Fig. 6e). Upon treatment, Hippo-strong cells exhibited reduced spreading and formed compact aggregates, whereas the morphology of Hippo-weak cells remained unchanged, consistent with the siRNA experiments (Fig. 6f).

Finally, to enable clinical application, we sought a tissue-based biomarker for stratifying patients with the Hippo-strong state. To prioritize candidates compatible with IHC-based diagnostics, we restricted our analysis to non-secretory proteins and calculated the AUROC score for each gene in the aligned TCGA expression dataset. *NNMT* (Nicotinamide N-methyltransferase), a cytoplasmic enzyme, emerged as the top-ranked candidate, distinguishing Hippo-strong tumors with high accuracy (AUROC = 0.916; Fig. 6g, h, S9d). Although the mechanistic link between the Hippo pathway and *NNMT* remains to be elucidated, high *NNMT* expression correlated with increased sensitivity to both ITGAV knockout and Cilengitide treatment (Fig. 6i, j), supporting its utility as a stratification marker. Consistent with this, *NNMT* is highly expressed across many cancer types, including kidney and pancreatic cancers, and elevated NNMT expression has been independently associated with poor overall survival ^50^ — matching the poor outcomes observed in Hippo-strong tumors (Fig. 6k). Collectively, these findings nominate NNMT as a candidate diagnostic biomarker to identify patients with aggressive Hippo-strong cancers who may benefit from integrin αVβ5-targeted therapies (Fig. 6l).

## Discussion

In this study, we introduced a pathway-centric dimensionality reduction framework to map the functional diversity across cancer cell lines. By integrating gene expression and CRISPR-based knockout effect data from DepMap, our analysis revealed five distinct functional states of the Hippo pathway. A comparative analysis using both computational and experimental approaches revealed that cells in the ‘Hippo-strong’ cluster exhibit enhanced cell-cell adhesion and a corresponding vulnerability to integrin αV inhibition. To translate these findings to a clinical context, we applied machine learning to tumor datasets and identified *NNMT* expression as a potential biomarker of the Hippo-strong signature, highlighting a novel therapeutic opportunity for a subset of cancers.

Our analysis provides a mechanistic framework for the ‘Hippo-strong’ phenotype, which is characterized by marked growth upregulation following knockout of Hippo pathway genes (Fig. 2). Based on our findings, we propose three interrelated feedback loops that sustain this ‘Hippo-strong’ state. The first is a positive feedback loop between YAP/TAZ and the extracellular matrix (ECM). We observed that Hippo-strong cells concurrently exhibit high baseline YAP/TAZ activity and high expression of ECM components (Figs. 2, 3). This is likely driven by an established feedback circuit where YAP/TAZ upregulates the ECM, and a stiffened ECM, in turn, promotes further YAP/TAZ activity ^51^. Second, the Hippo pathway establishes another positive feedback loop by enhancing cell-cell adhesion independently of YAP/TAZ (Figs. 3-5). This elevated adhesion plausibly enforces contact inhibition, which further activates the Hippo pathway ^52^. The third mechanism is a negative feedback loop. In Hippo-strong cells, several Hippo pathway components including *AMOTL2, KIRREL1, NRP1, LATS2*, and *PTPN14*, are highly expressed (Fig. S4b). The expression levels of these genes exhibit a strong correlation with YAP/TAZ targets such as *CCN1* and *CCN2* across cancer cell lines (Fig. S4c), suggesting that YAP/TAZ upregulates these Hippo pathway components to inhibit itself. Indeed, *LATS2, PTPN14,* and *AMOTL2* are reported to be YAP/TAZ downstream targets ^20^. Collectively, the concurrent upregulation of Hippo components, YAP/TAZ targets, and ECM genes, together with these intertwined positive and negative feedback circuits, possibly maintains the distinctive Hippo-strong phenotype.

Our analysis suggested that Cilengitide is a potential drug for Hippo-strong cells (Fig. 6). Notably, Cilengitide reached clinical trials against several cancers, including phase III clinical trials for glioblastoma, but failed to show the survival improvement and yet be approved ^53–56^. This failure has been proposed to result from alternative integrins supported by the internal extracellular matrix, which may mitigate the drug’s effects ^57^. Another possible explanation for clinical failure is the discrepancy between cell-line models and tumor samples. Glioblastoma and thyroid cancer cell lines are known to diverge markedly from their parental tumors ^45^. Importantly, our machine learning–based transfer analysis revealed that the fraction of Hippo-strong in glioblastoma is substantially lower in tumors than DepMap cell lines (Fig. S9c), potentially explaining Cilengitide’s limited efficacy in glioblastoma trials. Thus, these findings suggest that Cilengitide may warrant preclinical re-evaluation in Hippo-strong/NNMT-high tumor models, such as patient-derived xenografts or patient-derived organoids.

Conceptually, the dimensionality reduction approach of StateMap relates to supervised PCA (SPCA) ^58^, as both methods pre-select features based on their relevance to a target variable (in this case, CRISPR knockout fitness data) prior to projecting them into a lower-dimensional space. StateMap differs in two respects: feature selection uses metrics tightly linked to biological function (co-dependency modules and mutual information) rather than univariate correlation, and TriMAP is used in place of linear PCA to preserve both local and global non-linear structure. These choices extend the core concept of SPCA to map the heterogeneous pathway states across cancer cell lines. Nevertheless, two methodological limitations remain. First, feature selection was not fully automated and did not rely on fixed universal thresholds, limiting scalability (Fig. 1) ^59^. Second, because the selected feature set was small, knockout effects and expression levels were integrated by simple concatenation and normalization; broader applications involving many gene features or additional omics layers will require more scalable data-integration strategies ^60–62^, such as canonical correlation analysis (CCA), Similarity Network Fusion, or Multi-Omics Factor Analysis (MOFA) ^61,63,64^. We therefore view automated feature-selection and data-integration frameworks as future extensions of StateMap.

## Supplementary Figure legends

**Figure S1**

**(a)** Histograms showing the distribution of cell lines based on their Chronos scores for representative genes. E(X = 0.05) and E(X = 0.95) indicate the 5th and 95th percentile scores, respectively, for each gene. **(b)** Scatter plot showing the relationship between E(X = 0.05) and E(X = 0.95) across all genes. The red curve represents a smoothing spline function that predicts E(X = 0.95) based on E(X = 0.05). The vertical deviation from this prediction curve is defined as “selectivity.” **(c)** Q–Q plot comparing the selectivity distribution with a normal distribution. The red and orange lines indicate the regression and +3 sigma thresholds, respectively. **(d)** Cluster map showing the pairwise correlation of Chronos scores for the 129 most selective genes (>3 sigma) across cell lines. **(e)** Cluster map showing the pairwise correlation of Chronos scores for the Hippo pathway, TP53 pathway, mTOR negative regulators, and SAGA complex genes. **(f)** Scatter plot showing the relationship between 1D Hippo activity scores and WWTR1 expression levels. **(g)** Scatter plot showing the relationship between 1D Hippo activity scores and CCN1 expression levels. **(h)** Scatter plot showing the relationship between 1D Hippo activity scores and CCN2 expression levels.

**Figure S2**

**(a)** Dimensionality-reduction embeddings of Hippo gene data generated using PCA, UMAP, and t-SNE. **(b)** Global scores of the embeddings in (a). **(c)** Ranked gene scatter plot of mutual information scores between gene expression levels and Hippo pathway gene knockout effects. **(d)** Dimensionality-reduction embeddings of Hippo gene data from Chronos data (KEGG, hsa04390, Hippo signaling activity) and expression data (combined gene sets from Kanai et al.^21^ and Wang et al. ^20^) generated using TriMAP. **(e)** Graph-Laplacian smoothness penalty (k-NN, k=30) for the 1D Hippo pathway activity scores across the embeddings. Corr: correlation; Lit: literature (combined gene sets from Kanai et al.^21^ and Wang et al.^20^). **(f)** Tables showing the top mutual information (MI) genes for the TP53, mTOR, and SAGA complex pathways. For TP53 targets, the census p53 target genes were used ^26^. For mTOR and SAGA, such prior information was unavailable, so the top-ranking genes were directly selected for StateMap.

**Figure S3**

**(a)** Line plot showing the eigenvalues across different values of *k* in spectral clustering for the original concatenated feature matrix before dimensionality reduction. **(b)** Line plot showing the eigenvalues across different values of *k* in spectral clustering for the 2D embeddings shown in Figure 1d. **(c)** Line plot showing the eigengap after index *k* corresponding to the eigenvalues in (b). **(d)** Line plot showing the corresponding eigengap after index *k* from (c). **(e)** Consensus matrix for the 2D embeddings at *k* = 5. The clustering was performed 1,000 times, with 10% of the samples randomly masked in each iteration. **(f)** Line plot showing the PAC (Proportion of Ambiguously Clustered) scores for *k* = 2 to 10.

**Figure S4**

**(a)** Heatmap showing expression levels of 22 YAP/TAZ target genes ^20^. **(b)** Heatmap showing the expression levels of 11 Hippo pathway genes used in StateMap construction. **(c)** Bar plot showing the correlation between the expression levels of the genes in (b) and that of CCN1.

**Figure S5**

**(a)** Distribution of Hippo-derived clusters across different cell types. **(b)** Pie chart showing the distribution of cancer types within each cluster. **(c)** Violin plots showing the distribution of expression levels across the five clusters for the indicated neuroendocrine-related genes. *P* values were calculated using a two-tailed Mann–Whitney *U* test. \*\*\**P*□<□0.001. **(d)** Violin plots showing the distribution of expression levels across the five clusters for the indicated genes involved in noncanonical adhesion.

**Figure S6**

Stacked bar chart showing the distribution of the five clusters across subtypes.

**Figure S7**

**(a)** Bar charts showing expression levels of ITGAV and ITGB5 in the cell lines used in Fig. 4. **(b)** Schematic illustration of CRISPR-Cas9 mediated deletion knockout of TAOK1. **(c)** PCR-based validation of TAOK1 knockout in RPE1 cells. **(d)** Sanger sequencing of the genomic region in TAOK1 KO cells. **(e)** Western blot showing depletion efficiency of ITGAV and cleavage of PARP, an apoptosis marker. **(f)** Time-lapse images of RPE1-Fucci cells treated with a CDK4/6 inhibitor and the indicated siRNAs.

**Figure S8**

**(a)** Gene Ontology (GO) enrichment analysis of genes upregulated in TAOK1 knockout (KO) cells compared to wild-type (WT) cells. **(b)** Heatmaps showing expression levels of upregulated genes associated with the GO term “antigen processing and presentation of exogenous peptide antigen via MHC class II” in triplicate WT and KO samples (left), and across the five Hippo-derived clusters (right).

**Figure S9**

**(a)** Bar chart showing the balanced accuracy for the same three models. **(d)** Accumulated bar chart showing the distribution of true versus predicted labels across all clusters. **(c)** Scatter plot comparing the proportion of Hippo-strong cells in DepMap cell lines and tumor samples. **(d)** ROC curve for *NNMT* expression.

## Material and Methods

### Cell culture

The cell lines used in this study were obtained from various sources. hTERT RPE1 cells were purchased from ATCC (American Type Culture Collection). RPE1 Fucci cells were a gift from David Pellman^66^. U2OS cells were purchased from ECACC (European Collection of Authenticated Cell Cultures). PANC1, K562 and MCF7 cells were obtained from RIKEN BRC (RIKEN BioResource Research Center). YKG1, KMRC1, T24, KYSE270, and HEC251 cells were obtained from JCRB (Japanese Cancer Research Resources Bank).

hTERT RPE1 cells were cultured in DMEM/Ham’s F12 supplemented with 10% heat-inactivated FBS and 1% penicillin-streptomycin. KYSE270 cells were cultured in RPMI-1640/Ham’s F12 supplemented with 2% heat-inactivated FBS and 1% penicillin-streptomycin. U2OS, DLD1, and MCF7 cells were cultured in DMEM/High Glucose supplemented with 10% heat-inactivated FBS and 1% penicillin-streptomycin. YKG1 and KMRC1 cells were cultured in DMEM/Low Glucose supplemented with 10% heat-inactivated FBS and 1% penicillin-streptomycin. PANC1 cells were cultured in RPMI-1640 with 10% heat-inactivated FBS and 1% penicillin-streptomycin. K562 cells were cultured in IMDM with 10% heat-inactivated FBS and 1% penicillin-streptomycin. HEC251 cells were cultured in EMEM with 15% heat-inactivated FBS, 1% L-glutamine, and 1% penicillin-streptomycin. T24 cells were cultured in EMEM with 10% heat-inactivated FBS, 1% L-glutamine and 1% penicillin-streptomycin. All cells were maintained at 37°C in a humidified incubator with 5% CO₂.

### siRNA-mediated gene knockdown

siRNAs were transfected at a final concentration of 10 nM using Lipofectamine RNAiMAX (2 µL/well; Thermo Fisher Scientific). For immunofluorescence staining and Western blotting experiments, siRNA transfection was conducted 24 hours after cell seeding, followed by incubation for 24 to 48 hours before fixation or protein extraction. For live-cell imaging experiments, imaging was performed from the time of siRNA transfection up to 48 to 60 hours. The siRNAs used in this study were purchased from Thermo Fisher Scientific and included the following: TAOK1 (s33291), KIRREL1 (s30530), NF2 (s194648), RNF146 (s37822), PTPN14 (s11531), YAP1 (s20367), WWTR1 (s24787), ITGAV (s7568), and a negative control siRNA (4390843).

### Reagents and Chemicals

The following drugs and reagents were used at the indicated final concentrations: Palbociclib (CDK4/6 inhibitor, 300 nM; Selleck Chemicals LLC, S1116), Doxorubicin (1 μM; Toronto Research Chemicals Inc., TRC-D558000), Cilengitide (0.03 - 30 μM, Selleck, S6387), and YAP/TAZ inhibitor-1 (100 nM; MedChemExpress, HY-111429). For live-cell imaging, the fluorescent probes SiR-DNA (100 nM; Cytoskeleton Inc., CY-SC007) and SiR-Actin (100 nM; Cytoskeleton Inc., CY-SC001) were used to label nuclei and the actin, respectively.

### Generation of knockout (KO) cell lines

RPE1-TAOK1 KO cells were generated by excising the entire TAOK1 gene using a dual-site CRISPR-Cas9 approach, following the CRISPR-del method ^44^. Single-guide RNAs (sgRNAs) were designed to target the first intron (5’-GUGUACCUCUGAAUCCCACG-3’) and the last exon (5’-GCAAGUGGCCCUAAUAUACC-3’) of the *TAOK1* locus. These sgRNAs were synthesized by in vitro transcription of DNA oligonucleotide templates using the HiScribe T7 Transcription Kit (New England Biolabs) and subsequently purified with the RNA Clean & Concentrator kit (ZYMO RESEARCH). Purified sgRNAs and Alt-R® S.p. HiFi Cas9 Nuclease V3 (IDT, 1081061) were introduced into wild-type RPE1 cells by electroporation. After several days of culture, single-cell clones were isolated in 96-well plates. Successful knockout was validated in these clones by Western blotting (Fig. 4f), Sanger sequencing (Fig. S7d), and PCR (Fig. S7c) using the following primers: WT detection Fw (5’-GCTCAGGGGCACCAAATACT-3’), WT detection Rv (5’-AGACACGTGCCTATTGGTGG-3’), KO detection Fw (5’-ACTTCTTCTTTGCATAGGAGTCTTT-3’), and KO detection Rv (5’-TTGGAGGGGCACTCCTAAGT-3’).

### Immunofluorescence

Cells were prepared for imaging on either 12-mm or 15-mm coverslips (for adherent cells) or collected onto filters via centrifugation (for suspension cells), as detailed in ^67^. The Cells were then fixed with 4% paraformaldehyde (Nacalai Tesque, 09154-85) for 15 minutes. Cells were permeabilized in PBS-X (0.05% Triton X-100 in PBS) for 5 minutes, followed by blocking with 1% BSA in PBS-X (blocking buffer) for 15 minutes. All fixation and staining steps were performed at room temperature. Following blocking, cells were incubated for 2 hours with rabbit primary antibodies against YAP1 (Cell Signaling Technology, 14074) or WWTR1 (Cell Signaling Technology, 4883). After three washes in PBS, the cells were incubated for 1 hour with an Alexa Fluor 488-conjugated donkey anti-rabbit IgG secondary antibody (Thermo Fisher Scientific, A32790). To visualize F-actin and nuclei, cells were stained with Phalloidin-iFluor 555 (1:500 or 1:1,000; Cayman Chemical, 20552) for 30 minutes and Hoechst 33258 (1:5,000; Nacalai Tesque, 19173-41) for 5 minutes. All incubations were conducted at room temperature. Finally, coverslips were washed three times with PBS and mounted onto slides using a mounting medium. Images were acquired on an Axioplan2 fluorescence microscope or a ZEISS LSM 980 with Airyscan 2 (Carl Zeiss).

### Live-cell imaging

Live-cell imaging was performed using a CQ1 Benchtop High-Content Analysis System (Yokogawa Electric) equipped with 10× and 20× objective lenses or a Cell Voyager CV1000 (Yokogawa Electric). Cells were seeded in 24-well glass-bottom SENSOPLATEs (Greiner Bio-One, 662892) for CQ1 imaging or in 35 mm 4-compartment glass-bottom dishes (Greiner Bio-One, 627870) for CV1000 imaging. Imaging was conducted at 37°C with 5% CO₂ at 15-minute intervals for 48 hours. SiR-DNA or SiR-Actin was added at a final concentration of 100 nM 3 hours before imaging. Acquired time-lapse images were analyzed using ImageJ.

### Cell aggregation index

The cell aggregation index was calculated from live-cell time-lapse images using ImageJ. First, each image was segmented into 200 μm × 200 μm subregions. Within each subregion, the total nuclear area was determined by image thresholding. The coefficient of variation (CV) of this nuclear area across all subregions was then calculated to quantify the degree of cell aggregation. A low CV indicated an even cell distribution, whereas a high CV signified aggregation due to the concentration of nuclei in a smaller number of subregions.

### Western blotting

Cells were lysed using lysis buffer containing 20 mM Tris-HCl (pH 7.5), 50 mM NaCl, 1% Triton X-100, 5 mM EGTA, 1 mM DTT, 2 mM MgCl₂, and a 1:1000 dilution of protease inhibitor cocktail (Nacalai Tesque, 25955-11). SDS sample buffer (Nacalai Tesque, 09499-14) was added to each sample, followed by boiling at 95°C for 5 minutes and membrane disruption by ultrasonication. Proteins were then separated by SDS-PAGE and transferred onto an Immobilon-P membrane (Merck Millipore, IPVH00010) using a semi-dry transfer method. After transfer, the membrane was blocked with 5% skim milk in PBS-T (0.02% Tween in PBS) for 30 minutes, washed six times with PBS-T, and incubated overnight at 4°C with a primary antibody diluted in 5% BSA in PBS-T. The membrane was then washed six times with PBS-T, followed by incubation with a secondary antibody diluted in 5% skim milk in PBS-T at room temperature for 1 hour, and washed again six times with PBS-T. For protein detection, Chemi-Lumi One L, Super, or Ultra reagents (Nacalai Tesque, 07880, 02230, 11644) were used. Band images were acquired using ChemiDoc XPS+ and Image Lab Software (Bio-Rad).

For Western blotting, the following primary antibodies were used. Antibodies raised in rabbit (all used at 1:1,000 dilution) were: anti-ITGAV (Abcam, ab179475), anti-PARP (Cell Signaling Technology, 9542), anti-TAOK1 (Proteintech, 26250-1-AP), anti-YAP1 (Cell Signaling Technology, 14074), and anti-WWTR1 (Cell Signaling Technology, 4883). Antibodies raised in mouse (both used at 1:1,000 dilution) were: anti-HSP90 (BD Biosciences, 610419) and anti-α-Tubulin (Sigma-Aldrich, T5168). Secondary antibodies were HRP-conjugated Goat Anti-Mouse IgG (H+L) (1:10,000; Promega, W4021) and HRP-conjugated Goat Anti-Rabbit IgG (H+L) (1:10,000; Promega, W4011).

### MTT assay

Cells were seeded at a density of 1L×L10^4^ cells/mL in a 96-well plate. The medium was replaced with fresh medium containing various concentrations of cilengitide. The plate was then incubated for 3 days (for U2OS, YKG-1, KMRC-1, T-24, KYSE270 cell lines) or 6 days (for HEC-251 cell line). Following incubation, 10 μL of MTT solution (ThermoFisher, 5 mg/mL in PBS) was added to each well. Subsequently, the medium was discarded, and MTT formazan crystals were dissolved by adding isopropanol. The optical density was measured at 600 nm using a FLUOstar OPTIMA microplate reader (BMG LABTECH). Cell viability was calculated by normalizing the optical density readings to those of the control treatment.

### Genome-wide CRISPR screening

A genome-wide CRISPR-Cas9 screen was performed using hTERT-RPE1 cells, which serve as a model for the ‘Hippo-strong’ cluster. To achieve this, hTERT-RPE1 cells stably expressing Cas9 were created via lentiviral infection. For the lentiviral infection, the plasmid pKLV2-EF1a-BsdCas9-W (Addgene #67978) was introduced into 293FT cells along with psPAX2 (Addgene #12260) and pCMV-VSV-G (Addgene #8454). The resulting recombinant lentiviruses were then used to infect hTERT-RPE1 cells, and single cells were isolated through limited dilution. To enable the selection of lentivirus sgRNA library transduced cells using puromycin, we knocked out the puromycin acetyltransferase (PAC) gene, which is expressed in hTERT-RPE-1 cells stably expressing Cas9. An sgRNA targeting PAC (5′-TGTCGAGCCCGACGCGCGTG-3′, ^68^), was synthesized in vitro. This sgRNA was then transfected into hTERT-RPE1 cells that stably expressed the Cas9 protein. Single cells were subsequently isolated into 96-well plates using cellenONE (Cellenion) according to the manufacturer’s instructions. Subsequently, the screening was performed by using a lentiviral library (The Sabatini/Lander CRISPR pooled library, Addgene#1000000095, ^43^) targeting protein-coding genes across the genome. Cells were transduced at a low multiplicity of infection (MOI = 0.3) and subsequently selected with puromycin (3 µg/mL) to eliminate non-transduced cells. From 12 to 27 days post-transduction, samples were collected every three days. Genomic DNA was extracted using NucleoSpin^®^ Blood XL (MACHEREY-NAGEL), and sgRNA sequences were amplified by genomic PCR and subjected to next-generation sequencing (NGS). Gene knockout effects were evaluated based on sgRNA counts collected at six time points between day 12 and day 27 using the Chronos algorithm (https://github.com/broadinstitute/chronos) ^69^. Raw Chronos dependency values were scaled such that the mean of the Achilles common essential control genes was set to –1, and the mean of the nonessential control genes was set to 0.

### RNA sequencing

RPE1 wild-type and TAOK1-knockout samples (three biological replicates each) were submitted to KOTAI Bio Inc. for whole-genome transcriptome analysis. A count table was produced using the nf-core rnaseq workflow (https://zenodo.org/records/16892755). Reads were mapped with STAR^70^ and assigned to genes/transcripts using RSEM ^71^. Differential gene expression analysis was performed with edgeR ^72^.

### Webster Analysis for 1D Pathway Scores

To derive one-dimensional pathway activity scores, we utilized the Webster framework to decompose the dependency matrix^16^. Webster was applied to the DepMap 26Q1 dataset to extract latent functions using the parameters established in the original study (220 functions, with each gene attributed to 4 functions) ^73^. The dictionary values corresponding to specific latent factors (e.g., V13 for the Hippo pathway, V30 for the SAGA complex, V46 for the TP53 pathway, and V216 for mTOR negative regulators) were extracted and utilized as 1D pathway activity scores for comparative analyses across cell lines.

### Calculation of Gene Selectivity

Gene selectivity was calculated to quantify the variation of a gene’s dependency across different cell lines. Our method builds upon the approach described by Shimada et al. (2021) ^74^, with two key differences: (1) we used only CRISPR gene effect scores, whereas the original study integrated both CRISPR and RNAi data; and (2) we employed a smoothing spline (smoothing factor = 100) to model the relationship between dependency scores, replacing the linear regression used in the original study (Fig. S1a-c).

For each gene, the bottom 5% (E0.05) and top 5% (E0.95) of gene effect scores were extracted. A smoothing spline function, f(E0.05), was fitted to predict E0.95. Finally, gene selectivity was calculated as:

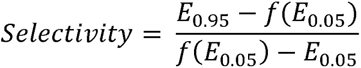

### Dimensionality reduction

Gene knockout effect data from CRISPR screening (Chronos score) and gene expression data from RNA sequencing were obtained from the DepMap 22Q2 dataset^75^. Cell lines present in only one dataset or with missing values were excluded, resulting in gene effect data for 17,386 genes and expression data for 19,221 genes across 1,001 cell lines.

To select Hippo-related genes for dimensionality reduction, hierarchical clustering was performed on Chronos scores of the top 129 selective genes (>3 sigma from the regression line). A large cluster containing 11 Hippo-related genes was identified and used for subsequent analyses (Fig. S1d). These genes were RNF146, LATS2, KIRREL1, AMOTL2, NF2, PDCD10, FRYL, NRP1, PTPN14, TAOK1, and MAP4K4 (Fig. 2a). Except for FRYL, all of these genes have been previously reported to be associated with the Hippo pathway ^76–81^. The role of FRYL in the Hippo pathway is not well established. However, its paralog FRY has been reported to regulate YAP and TAZ in conjunction with NDR1^82^, and FRYL was therefore included in our analysis.

To identify gene expression profiles most closely linked to functional dependencies, we calculated mutual information (MI) between the knockout effects of pathway-core genes and genome-wide gene expression levels. The Chronos scores of the 11 Hippo-related genes were normalized, and MI was computed between their mean score and the expression level of each gene. MI scores were calculated using the mutual_info_regression function from the scikit-learn Python library with default parameters. Genes with the highest MI scores were selected as expression features for StateMap integration. For the Hippo and TP53 pathways, previously curated gene^20,21,26^ were used as prior knowledge in combination with MI scores, whereas for the mTOR and SAGA complexes, the top-ranking MI genes were selected directly.

Chronos scores of the 11 Hippo-related genes, together with the expression of YAP1, WWTR1, CCN1, and AXL, were standardized and embedded into two dimensions using TriMAP^22^ (https://github.com/eamid/trimap). TriMAP was run with the parameters n_outliers=10 and n_inliers=10, while all other settings were left at their default values.

### Spectral Clustering and Evaluation

To objectively determine the optimal number of functional states, we performed the spectral clustering on the low-dimensional embedding coordinates generated by StateMap^28^. A symmetric k-nearest-neighbor (kNN) distance graph was constructed with k = 30.

Distances were converted into an affinity matrix using a radial basis function (RBF) kernel, with the bandwidth parameter, sigma, set to the median of the nonzero distances. The normalized graph Laplacian of this affinity matrix was then computed, and the 15 smallest eigenvalues were extracted. The optimal number of clusters was determined by identifying the maximum eigengap. Based on a local maximum in the eigenvalue gap, k = 5 was selected ^29^.

To assess the stability of this selection, consensus clustering was performed ^30^. The dataset was repeatedly subsampled 1,000 times, with 90% of samples randomly retained and 10% masked in each iteration. For each subsample, the kNN affinity graph reconstruction and spectral clustering steps were repeated. Clustering robustness was quantitatively evaluated by constructing a consensus matrix and calculating the proportion of ambiguously clustered pairs (PAC) score across a range of cluster numbers, k = 2 to 10^31^. The PAC score was defined as the fraction of sample pairs with pairwise consensus values between 0.1 and 0.9. Because k = 5 yielded a locally minimal PAC score and was consistent with the eigengap analysis, it was selected as the final number of clusters. All clustering steps were implemented using the scikit-learn and SciPy Python libraries.

### Smoothness

For each candidate embedding, smoothness of the Hippo pathway signal was quantified by a local-variance score over a k-nearest-neighbor graph (k = 15, Euclidean distance) built in the embedding space. Edge weights were computed from a Gaussian kernel on squared neighbor distances and row-normalized so that the weights from each cell to its neighbors sum to one. For every cell, the weighted sum of squared signal differences between that cell and its neighbors was computed, and the global score was taken as the mean across cells. This score is related to standard graph-Laplacian-based smoothness measures used in manifold learning, with lower values indicating that the signal varies more gradually along the embedding’s neighborhood structure.

### EMT score

E-score and M-score were calculated following the method described previously ^37^. For each cell line, expression levels (log₂TPM) were normalized to have a mean of 0 and a standard deviation of 1. The averaged expression levels of pan-cancer epithelial and mesenchymal marker genes described ^36^ were then calculated, representing the E-score and M-score, respectively.

### Machine learning classification for TCGA data

Machine learning models predicting Hippo state clusters were trained on DepMap expression data aligned with tumor samples using Celligner ^45^. This approach followed a previous study that used machine learning to predict gene perturbation effects in TCGA samples ^46^. To create a high-confidence training set, a non-linear Support Vector Machine (SVM) classifier was first trained on the 2D TriMap coordinates to predict the five clusters. Only samples with a prediction probability of 0.9 or higher were selected as “far from boundary” samples. The main gene expression data was then filtered to include only these high-confidence samples. A feature selection process was then applied to the gene expression data. An ANOVA F-test was performed to identify the top 30 most significant differentially expressed genes between all possible pairs of clusters. The final feature matrix used for training was the union of all selected genes from these pairwise comparisons, combined with the one-hot encoded lineage and subtype features.

Logistic regression, random forest, support vector machines, and XGBoost models were trained, and the hyperparameter was adjusted using Bayesian optimization to maximize balanced accuracy. To address class imbalance in the training data, all models were implemented within a pipeline utilizing SMOTETomek. Model performance was evaluated using 5-fold cross-validation. achieved comparable performance, with macro F1 scores and balanced accuracy values of approximately 0.7 and no significant differences (Fig. S9a, b). XGBoost with SMOTE was selected for subsequent analyses. Because Hippo-strong represented the smallest class, the decision threshold for this cluster was set at 0.2. TCGA patient survival data were obtained from the GDC Data Portal (https://portal.gdc.cancer.gov/).

### Statistical analysis

Statistical analyses were performed using Python. For two-group comparisons, either Student’s t-test or the Mann–Whitney U test was used. When Student’s t-test was applied, normality was assessed with the Shapiro–Wilk test and homogeneity of variances with Levene’s test. For multiple-group comparisons, one-way ANOVA followed by Dunnett’s post-hoc test was conducted, with normality and variance homogeneity evaluated in the same manner. Survival curves were estimated using the Kaplan–Meier method and compared with the log-rank test. A p-value < 0.05 was considered statistically significant.

## Code availability

All codes used to perform the analyses, generate figures, and implement the web application are available at GitHub (https://github.com/kkkkito/statemap).

## Acknowledgments

We gratefully acknowledge William C. Hahn and members of his laboratory, as well as members of the Kitagawa laboratory, for their constructive feedback. We also thank R. Anzai for implementing the web application on the server. This work was supported by JSPS KAKENHI grants (Grant numbers: 18K06246, 19H05651, 20K15987, 20K22701, 21H02623, 22H02629, 22K20624, 23K14176, 23H02627) from the Ministry of Education, Science, Sports and Culture of Japan, the PRESTO program (JPMJPR21EC) and the CREST program (JPMJCR22E1) of the Japan Science and Technology Agency, Takeda Science Foundation, The Uehara Memorial Foundation, The Research Foundation for Pharmaceutical Sciences, Koyanagi Zaidan, The Kanae Foundation for the Promotion of Medical Science, Kato Memorial Bioscience Foundation, Naito Foundation, Heiwa Nakajima Foundation, Sumitomo Foundation, Inamori foundation, and Tokyo Foundation for Pharmaceutical Sciences.

## Author contributions

K.K.I., T.H., T.C., and D.K. conceived the study.

K.K.I., T.H., T.C., and D.K. designed the study.

K.K.I. developed the web interface.

T.H., K.K.I., R.N., M.H., K. Kuroki, K. Kawai, H.U., S.G., S.S., and T.C. performed experiments.

T.H., K.K.I., T.C., and D.K. analyzed the data.

K.K.I., J.I., G.T., T.K., K.A., and Y.S. analyzed the CRISPR screen data.

K.K.I., wrote the original manuscript. T.C., and D.K. revised the manuscript.

All authors reviewed the manuscript.

## Declaration of generative AI and AI-assisted technologies in the writing process

During the preparation of this work, the authors used GPT4.1, GPT-4o, GPT-5, Gemini 2.5 Pro, Gemini 3 Pro, and Claude Opus 4.7 to improve language and readability. After using this service, the authors reviewed and edited the content as needed and take full responsibility for the content of the publication.

## References

1. Sanchez-Vega, F. et al. Oncogenic Signaling Pathways in The Cancer Genome Atlas. Cell 173, 321–337.e10 (2018).

2. Vogelstein, B., et al. Cancer Genome Landscapes. Science (1979). 339, 1546–1558 (2013).

3. Hanahan, D. & Weinberg, R. A. Hallmarks of Cancer: The Next Generation. Cell 144, 646–674 (2011).

4. Garraway, L. A. & Lander, E. S. Lessons from the Cancer Genome. Cell 153, 17–37 (2013).

5. Hänzelmann, S., Castelo, R. & Guinney, J. GSVA: Gene set variation analysis for microarray and RNA-Seq data. BMC Bioinformatics 14, (2013).

6. Barbie, D. A. et al. Systematic RNA interference reveals that oncogenic KRAS-driven cancers require TBK1. Nature 462, 108–112 (2009).

7. Schubert, M. et al. Perturbation-response genes reveal signaling footprints in cancer gene expression. Nat. Commun. 9, 20 (2018).

8. Szalai, B. & Saez-Rodriguez, J. Why do pathway methods work better than they should? FEBS Lett. 594, 4189–4200 (2020).

9. Vogel, C. & Marcotte, E. M. Insights into the regulation of protein abundance from proteomic and transcriptomic analyses. Nat. Rev. Genet. 13, 227–232 (2012).

10. Geffen, Y. et al. Pan-cancer analysis of post-translational modifications reveals shared patterns of protein regulation. Cell 186, 3945–3967.e26 (2023).

11. Liu, Y., Beyer, A. & Aebersold, R. On the Dependency of Cellular Protein Levels on mRNA Abundance. Cell 165, 535–550 (2016).

12. Tsherniak, A. et al. Defining a Cancer Dependency Map. Cell 170, 564–576.e16 (2017).

13. Dharia, N. V et al. A first-generation pediatric cancer dependency map. Nat. Genet. 53, 529–538 (2021).

14. Arafeh, R., Shibue, T., Dempster, J. M., Hahn, W. C. & Vazquez, F. The present and future of the Cancer Dependency Map. Nat. Rev. Cancer 25, 59–73 (2025).

15. Wainberg, M. et al. A genome-wide atlas of co-essential modules assigns function to uncharacterized genes. Nat. Genet. 53, 638–649 (2021).

16. Pan, J. et al. Sparse dictionary learning recovers pleiotropy from human cell fitness screens. Cell Syst. 13, 286–303.e10 (2022).

17. Shimada, K., Bachman, J. A., Muhlich, J. L. & Mitchison, T. J. shinyDepMap, a tool to identify targetable cancer genes and their functional connections from Cancer Dependency Map data. Elife 10, (2021).

18. Zheng, Y. & Pan, D. The Hippo Signaling Pathway in Development and Disease. Dev. Cell 50, 264–282 (2019).

19. Shannon, C. E. A Mathematical Theory of Communication. Bell System Technical Journal 27, 379–423 (1948).

20. Wang, Y. et al. Comprehensive Molecular Characterization of the Hippo Signaling Pathway in Cancer. Cell Rep. 25, 1304–1317.e5 (2018).

21. Kanai, R., Norton, E., Stern, P., Hynes, R. O. & Lamar, J. M. Identification of a Gene Signature That Predicts Dependence upon YAP/TAZ-TEAD. Cancers (Basel). 16, (2024).

22. Amid, E. & Warmuth, M. K. TriMap: Large-scale Dimensionality Reduction Using Triplets. 10.48550/arXiv.1910.00204 (2019) doi:10.48550/arXiv.1910.00204.

23. McInnes, L., Healy, J., Saul, N. & Großberger, L. UMAP: Uniform Manifold Approximation and Projection. J. Open Source Softw. 3, 861 (2018).

24. Van Der Maaten, L. & Hinton, G. Visualizing data using t-SNE. Journal of Machine Learning Research 9, 2579–2605 (2008).

25. Kanehisa, M., Furumichi, M., Tanabe, M., Sato, Y. & Morishima, K. KEGG: New perspectives on genomes, pathways, diseases and drugs. Nucleic Acids Res. 10.1093/nar/gkw1092 (2017) doi:10.1093/nar/gkw1092.

26. Fischer, M. Census and evaluation of p53 target genes. Oncogene 36, 3943–3956 (2017).

27. Oliner, J. D., Saiki, A. Y. & Caenepeel, S. The Role of MDM2 Amplification and Overexpression in Tumorigenesis. Cold Spring Harb. Perspect. Med. 6, a026336 (2016).

28. Ng, A. Y., Jordan, M. I. & Weiss, Y. On spectral clustering: Analysis and an algorithm. in Advances in Neural Information Processing Systems (2002).

29. von Luxburg, U. A tutorial on spectral clustering. Stat. Comput. 17, 395–416 (2007).

30. Monti, S., Tamayo, P., Mesirov, J. & Golub, T. Consensus Clustering: A Resampling-Based Method for Class Discovery and Visualization of Gene Expression Microarray Data. Mach. Learn. 52, 91–118 (2003).

31. Șenbabaoğlu, Y., Michailidis, G. & Li, J. Z. Critical limitations of consensus clustering in class discovery. Sci. Rep. 4, 6207 (2014).

32. Pearson, J. D. et al. Binary pan-cancer classes with distinct vulnerabilities defined by pro- or anti-cancer YAP/TEAD activity. Cancer Cell 39, 1115–1134.e12 (2021).

33. Sharma, P., Michaels, Y. S. & Pearson, J. D. Molecular basis and therapeutic implications of binary YAPOn/YAPOff cancer classes. Biochem. J. 482, 741–61 (2025).

34. Borromeo, M. D. et al. ASCL1 and NEUROD1 Reveal Heterogeneity in Pulmonary Neuroendocrine Tumors and Regulate Distinct Genetic Programs. Cell Rep. 16, 1259–1272 (2016).

35. Rodarte, K. E. et al. Neuroendocrine Differentiation in Prostate Cancer Requires ASCL1. Cancer Res. 84, 3522–3537 (2024).

36. Mak, M. P., et al. A Patient-Derived, Pan-Cancer EMT Signature Identifies Global Molecular Alterations and Immune Target Enrichment Following Epithelial-to-Mesenchymal Transition. Clinical Cancer Research 22, 609–620 (2016).

37. Pacini, C. et al. A comprehensive clinically informed map of dependencies in cancer cells and framework for target prioritization. Cancer Cell 42, 301–316.e9 (2024).

38. Legerstee, K. & Houtsmuller, A. A Layered View on Focal Adhesions. Biology (Basel). 10, 1189 (2021).

39. Pang, X. et al. Targeting integrin pathways: mechanisms and advances in therapy. Signal Transduct. Target. Ther. 8, 1 (2023).

40. Lock, J. G. et al. Clathrin-containing adhesion complexes. Journal of Cell Biology 218, 2086–2095 (2019).

41. Lukas, F. et al. Canonical and non-canonical integrin-based adhesions dynamically interconvert. Nat. Commun. 15, 2093 (2024).

42. Hakanpää, L. et al. Reticular adhesions are assembled at flat clathrin lattices and opposed by active integrin α5β1. Journal of Cell Biology 222, (2023).

43. Park, R. J. et al. A genome-wide CRISPR screen identifies a restricted set of HIV host dependency factors. Nat. Genet. 49, 193–203 (2017).

44. Komori, T. et al. A CRISPR-del-based pipeline for complete gene knockout in human diploid cells. J. Cell Sci. 136, (2023).

45. Warren, A. et al. Global computational alignment of tumor and cell line transcriptional profiles. Nat. Commun. 12, 22 (2021).

46. Shi, X. et al. Building a translational cancer dependency map for The Cancer Genome Atlas. Nat. Cancer 5, 1176–1194 (2024).

47. Dash, R. et al. Drug repurposing a compelling cancer strategy with bottomless opportunities. Indian J. Pharmacol. 55, 322–331 (2023).

48. Reardon, D. A., Nabors, L. B., Stupp, R. & Mikkelsen, T. Cilengitide: an integrin-targeting arginine–glycine–aspartic acid peptide with promising activity for glioblastoma multiforme. Expert Opin. Investig. Drugs 17, 1225–1235 (2008).

49. Dechantsreiter, M. A. et al. N-Methylated Cyclic RGD Peptides as Highly Active and Selective αVβ3 Integrin Antagonists. J. Med. Chem. 42, 3033–3040 (1999).

50. Wang, W., Yang, C., Wang, T. & Deng, H. Complex roles of nicotinamide N-methyltransferase in cancer progression. Cell Death Dis. 13, 267 (2022).

51. Zanconato, F., Cordenonsi, M. & Piccolo, S. YAP/TAZ at the Roots of Cancer. Cancer Cell 29, 783–803 (2016).

52. Gumbiner, B. M. & Kim, N.-G. The Hippo-YAP signaling pathway and contact inhibition of growth. J. Cell Sci. 127, 709–717 (2014).

53. Stupp, R. et al. Cilengitide combined with standard treatment for patients with newly diagnosed glioblastoma with methylated MGMT promoter (CENTRIC EORTC 26071-22072 study): a multicentre, randomised, open-label, phase 3 trial. Lancet Oncol. 15, 1100–1108 (2014).

54. Reardon, D. A. et al. Randomized Phase II Study of Cilengitide, an Integrin-Targeting Arginine-Glycine-Aspartic Acid Peptide, in Recurrent Glioblastoma Multiforme. Journal of Clinical Oncology 26, 5610–5617 (2008).

55. Alva, A. et al. Phase II study of Cilengitide (EMD 121974, NSC 707544) in patients with non-metastatic castration resistant prostate cancer, NCI-6735. A study by the DOD/PCF prostate cancer clinical trials consortium. Invest. New Drugs 30, 749–757 (2012).

56. Vansteenkiste, J. et al. Cilengitide combined with cetuximab and platinum-based chemotherapy as first-line treatment in advanced non-small-cell lung cancer (NSCLC) patients: results of an open-label, randomized, controlled phase II study (CERTO). Annals of Oncology 26, 1734–1740 (2015).

57. Bergonzini, C., Kroese, K., Zweemer, A. J. M. & Danen, E. H. J. Targeting Integrins for Cancer Therapy - Disappointments and Opportunities. Front. Cell Dev. Biol. 10, (2022).

58. Bair, E., Hastie, T., Paul, D. & Tibshirani, R. Prediction by Supervised Principal Components. J. Am. Stat. Assoc. 101, 119–137 (2006).

59. Saeys, Y., Inza, I. & Larrañaga, P. A review of feature selection techniques in bioinformatics. Bioinformatics 23, 2507–2517 (2007).

60. Ritchie, M. D., Holzinger, E. R., Li, R., Pendergrass, S. A. & Kim, D. Methods of integrating data to uncover genotype–phenotype interactions. Nat. Rev. Genet. 16, 85–97 (2015).

61. Huang, S., Chaudhary, K. & Garmire, L. X. More Is Better: Recent Progress in Multi-Omics Data Integration Methods. Front. Genet. 8, (2017).

62. Picard, M., Scott-Boyer, M.-P., Bodein, A., Périn, O. & Droit, A. Integration strategies of multi-omics data for machine learning analysis. Comput. Struct. Biotechnol. J. 19, 3735–3746 (2021).

63. Wang, B. et al. Similarity network fusion for aggregating data types on a genomic scale. Nat. Methods 11, 333–337 (2014).

64. Witten, D. M., Tibshirani, R. & Hastie, T. A penalized matrix decomposition, with applications to sparse principal components and canonical correlation analysis. Biostatistics 10, 515–534 (2009).

65. Zerbib, J. et al. Human aneuploid cells depend on the RAF/MEK/ERK pathway for overcoming increased DNA damage. Nat. Commun. 15, 7772 (2024).

66. Ganem, N. J. et al. Cytokinesis Failure Triggers Hippo Tumor Suppressor Pathway Activation. Cell 158, 833–848 (2014).

67. Miyazawa, M., Kawai, K., Yamamoto, S. & Kitagawa, D. Protocol for immunostaining of non-adherent cells and cellular structures using centrifugal filter devices. STAR Protoc. 6, 103934 (2025).

68. Lambrus, B. G. et al. A USP28–53BP1–p53–p21 signaling axis arrests growth after centrosome loss or prolonged mitosis. Journal of Cell Biology 214, 143–153 (2016).

69. Dempster, J. M. et al. Chronos: a cell population dynamics model of CRISPR experiments that improves inference of gene fitness effects. Genome Biol. 22, 343 (2021).

70. Dobin, A. et al. STAR: ultrafast universal RNA-seq aligner. Bioinformatics 29, 15–21 (2013).

71. Li, B. & Dewey, C. N. RSEM: accurate transcript quantification from RNA-Seq data with or without a reference genome. BMC Bioinformatics 12, 323 (2011).

72. Robinson, M. D., McCarthy, D. J. & Smyth, G. K. edgeRl1: a Bioconductor package for differential expression analysis of digital gene expression data. Bioinformatics 26, 139–140 (2010).

73. DepMap 26Q1 dataset.

74. Shimada, K., Bachman, J. A., Muhlich, J. L. & Mitchison, T. J. shinyDepMap, a tool to identify targetable cancer genes and their functional connections from Cancer Dependency Map data. Elife 10, 1–20 (2021).

75. Depmap. DepMap 22Q2 Public. figshare. Dataset. (2022).

76. Plouffe, S. W. et al. Characterization of Hippo Pathway Components by Gene Inactivation. Mol. Cell 64, 993–1008 (2016).

77. Paul, A. et al. Cell adhesion molecule KIRREL1 is a feedback regulator of Hippo signaling recruiting SAV1 to cell-cell contact sites. Nat. Commun. 13, 930 (2022).

78. Li, M. et al. NRP1 transduces mechanical stress inhibition via LATS1/YAP in hypertrophic scars. Cell Death Discov. 9, 341 (2023).

79. Wang, Y. et al. Stabilization of Motin family proteins in NF2-deficient cells prevents full activation of YAP/TAZ and rapid tumorigenesis. Cell Rep. 36, 109596 (2021).

80. Sun, B. et al. Programmed cell death 10 promotes metastasis and epithelial-mesenchymal transition of hepatocellular carcinoma via PP2Ac-mediated YAP activation. Cell Death Dis. 12, 849 (2021).

81. Michaloglou, C. et al. The Tyrosine Phosphatase PTPN14 Is a Negative Regulator of YAP Activity. PLoS One 8, e61916 (2013).

82. Irie, K., Nagai, T. & Mizuno, K. Furry protein suppresses nuclear localization of yes-associated protein (YAP) by activating NDR kinase and binding to YAP. Journal of Biological Chemistry 295, 3017–3028 (2020).

