## Supplemental figures for "Pathway-Centric Integration of CRISPR Fitness with Molecular Features Draws Cancer State Maps"

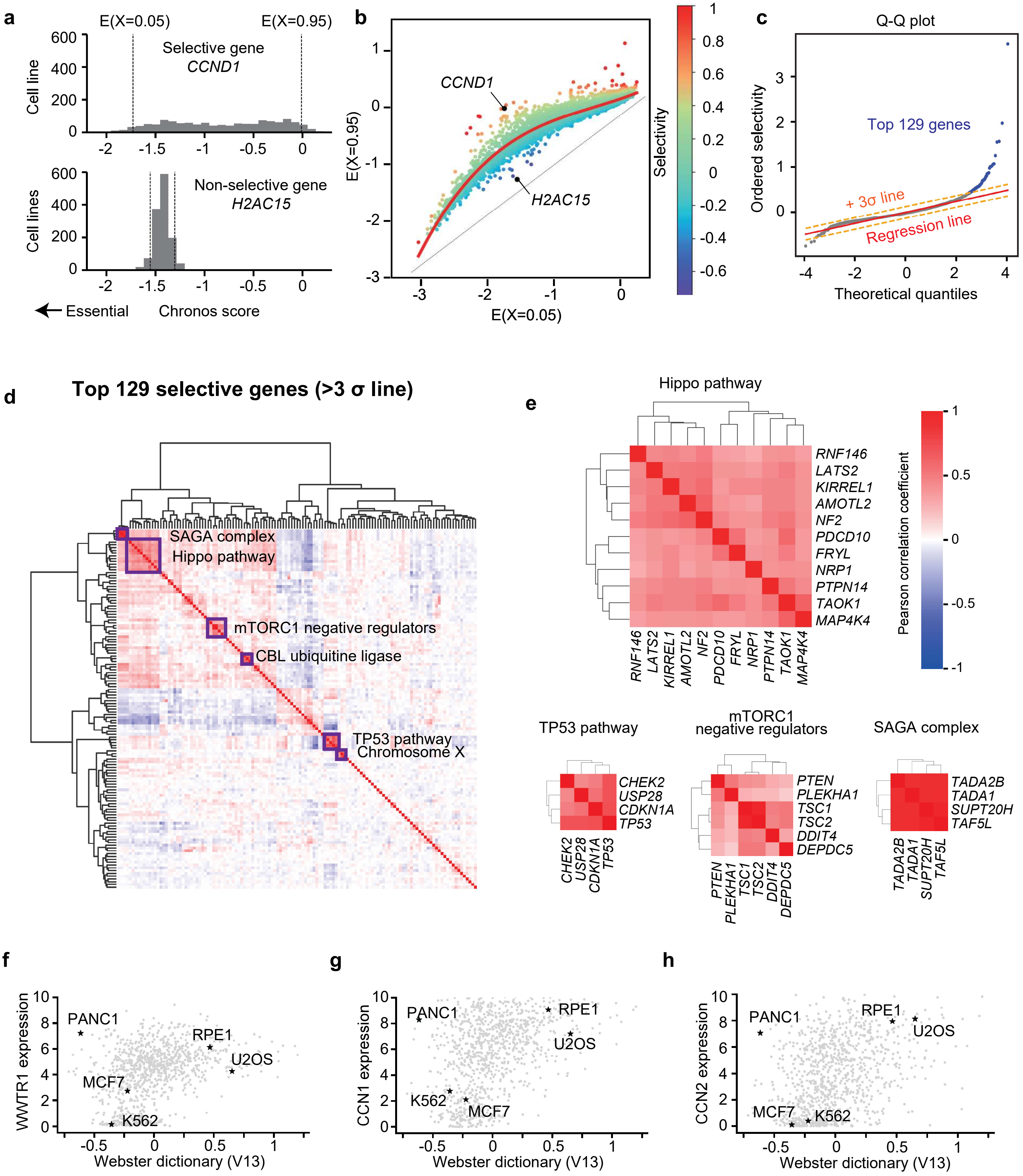

a
E(X=0.05)
E(X=0.95)
Selective gene
CCND1
Cell line
Non-selective gene
H2AC15
Cell lines
Essential
Chronos score
b
CCND1
H2AC15
E(X=0.95)
E(X=0.05)
Selectivity
c
Q-Q plot
Top 129 genes
+ 3σ line
Regression line
Ordered selectivity
Theoretical quantiles
d
Top 129 selective genes (>3 σ line)
SAGA complex
Hippo pathway
mTORC1 negative regulators
CBL ubiquitine ligase
TP53 pathway
Chromosome X
e
Hippo pathway
RNF146
LATS2
KIRREL1
AMOTL2
NF2
PDCD10
FRYL
NRP1
PTPN14
TAOK1
MAP4K4
Pearson correlation coefficient
TP53 pathway
CHEK2
USP28
CDKN1A
TP53
mTORC1 negative regulators
PTEN
PLEKHA1
TSC1
TSC2
DDIT4
DEPDC5
SAGA complex
TADA2B
TADA1
SUPT20H
TAF5L
f
WWTR1 expression
PANC1
RPE1
U2OS
MCF7
K562
Webster dictionary (V13)
g
CCN1 expression
PANC1
RPE1
U2OS
K562
MCF7
Webster dictionary (V13)
h
CCN2 expression
PANC1
RPE1
U2OS
MCF7
K562
Webster dictionary (V13)

a

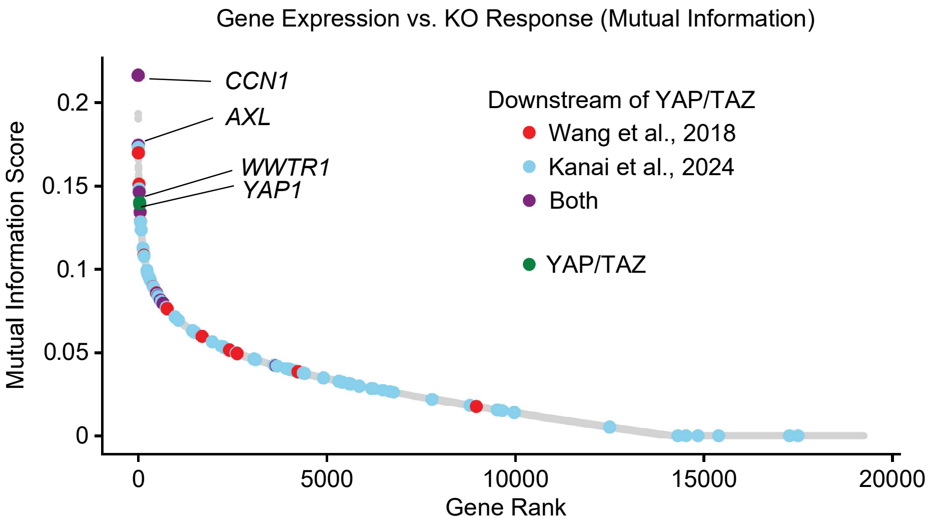


| Rank | Gene | MI | Wang et al., | Kanai et al., |
|---|---|---|---|---|
| 1 | *CCN1* † | 0.216 | x | x |
| 2 | *CAV1* | 0.193 | | |
| 3 | *TNFRSF12A* | 0.191 | | |
| 4 | *AXL* † | 0.175 | x | x |
| 5 | *FOSL1* | 0.173 | | x |
| 6 | *PXDC1* | 0.173 | | |
| 7 | *CCDC80* | 0.170 | x | |
| 8 | *UBTD1* | 0.169 | | |
| 9 | *ITGA3* | 0.168 | | |
| 10 | *GPX8* | 0.166 | | |
| 38 | *WWTR1* † | 0.140 | | |
| 39 | *YAP1* † | 0.139 | | |

† : Used in the Statemap

b c

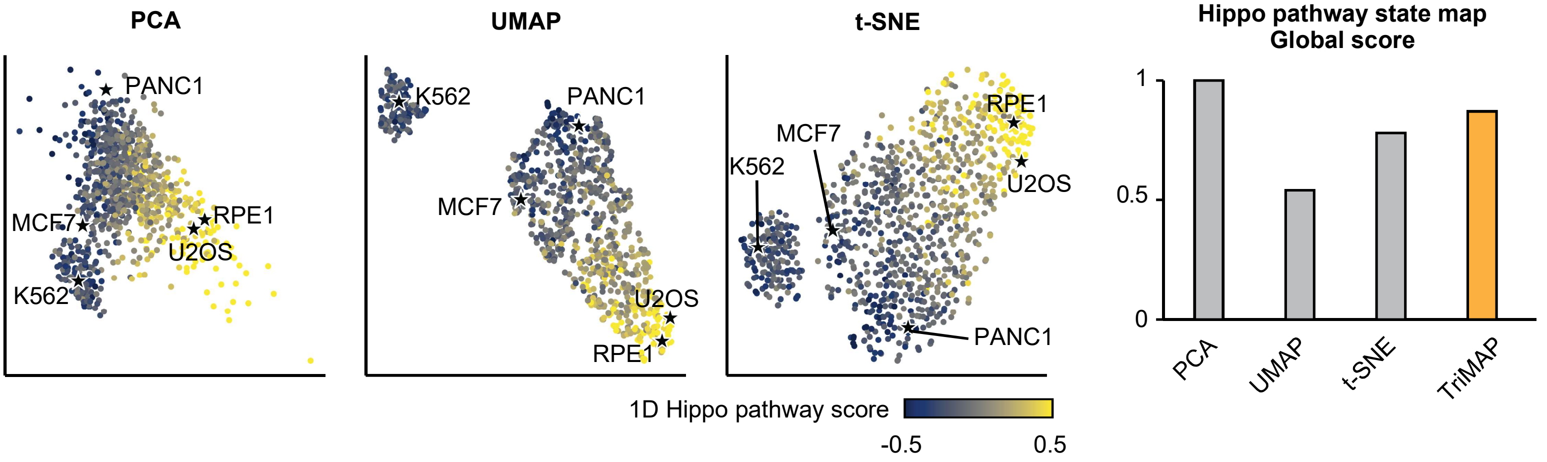


d

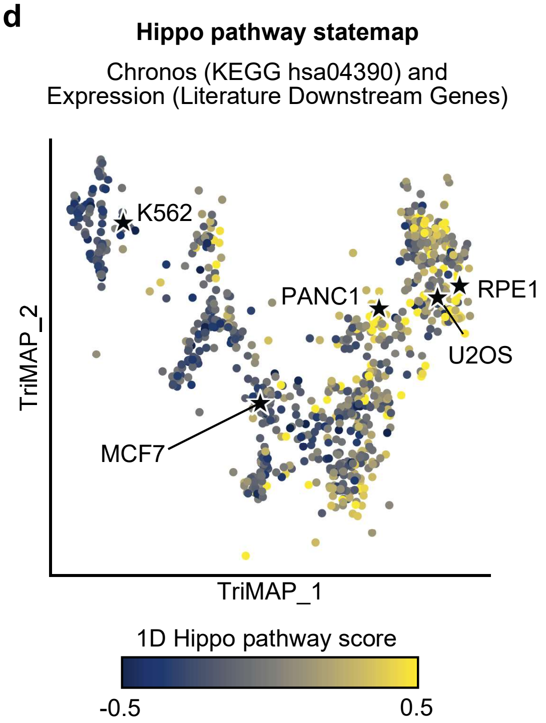


e

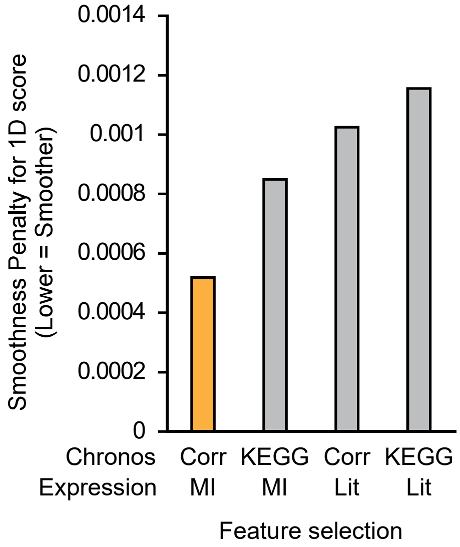


f

| | Rank | Gene | MI | Target |
|---|---|---|---|---|
| TP53 pathway | 1 | *CDKN1A* † | 0.241 | x |
| | 2 | *EDA2R* † | 0.191 | x |
| | 3 | *MDM2* † | 0.169 | x |
| | 4 | *PTCHD4* | 0.168 | |
| | 5 | *RPS27L* † | 0.156 | x |
| mTOR regulator | 1 | *CDC25C* † | 0.101 | |
| | 2 | *ZNF462* † | 0.101 | |
| | 3 | *ITSN1* | 0.100 | |
| | 4 | *SRD5A1* | 0.097 | |
| | 5 | *MAP4* | 0.097 | |
| SAGA complex | 1 | *GPX8* † | 0.171 | |
| | 2 | *PDGFC* † | 0.171 | |
| | 3 | *AXL* | 0.157 | |
| | 4 | *KIRREL1* | 0.153 | |
| | 5 | *AMOTL2* | 0.150 | |

† : Used in the Statemap

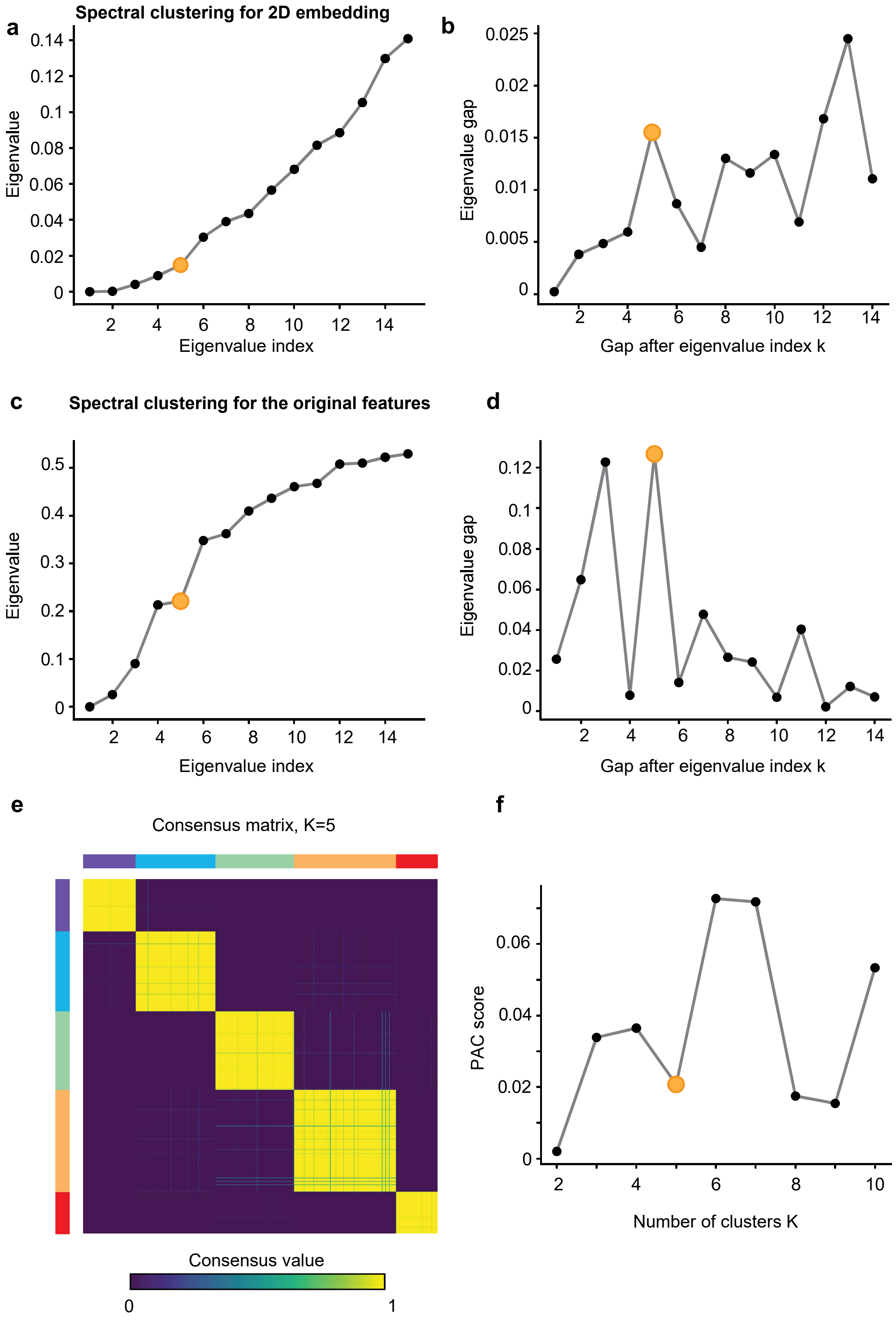

a
Spectral clustering for 2D embedding
Eigenvalue
Eigenvalue index
0
0.02
0.04
0.06
0.08
0.1
0.12
0.14
2
4
6
8
10
12
14
b
Eigenvalue gap
Gap after eigenvalue index k
0
0.005
0.01
0.015
0.02
0.025
c
Spectral clustering for the original features
Eigenvalue
Eigenvalue index
0
0.1
0.2
0.3
0.4
0.5
d
Eigenvalue gap
Gap after eigenvalue index k
0
0.02
0.04
0.06
0.08
0.1
0.12
e
Consensus matrix, K=5
Consensus value
0
1
f
PAC score
Number of clusters K
0
0.02
0.04
0.06

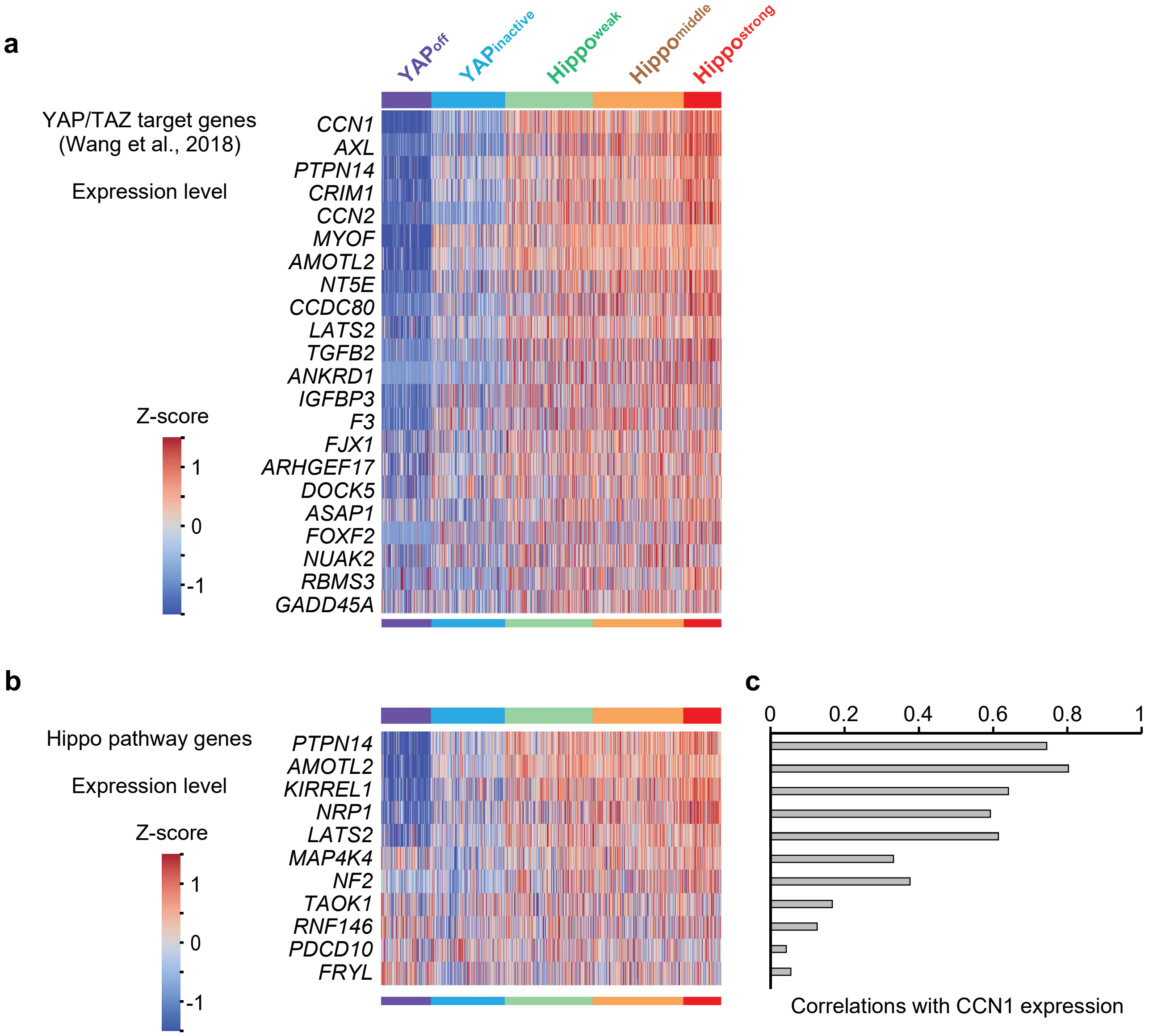
a
YAP^off
YAP^inactive
Hippo^weak
Hippo^middle
Hippo^strong
YAP/TAZ target genes
(Wang et al., 2018)
Expression level
Z-score
1
0
-1
CCN1
AXL
PTPN14
CRIM1
CCN2
MYOF
AMOTL2
NT5E
CCDC80
LATS2
TGFB2
ANKRD1
IGFBP3
F3
FJX1
ARHGEF17
DOCK5
ASAP1
FOXF2
NUAK2
RBMS3
GADD45A
b
Hippo pathway genes
Expression level
Z-score
1
0
-1
PTPN14
AMOTL2
KIRREL1
NRP1
LATS2
MAP4K4
NF2
TAOK1
RNF146
PDCD10
FRYL
c
0
0.2
0.4
0.6
0.8
1
Correlations with CCN1 expression

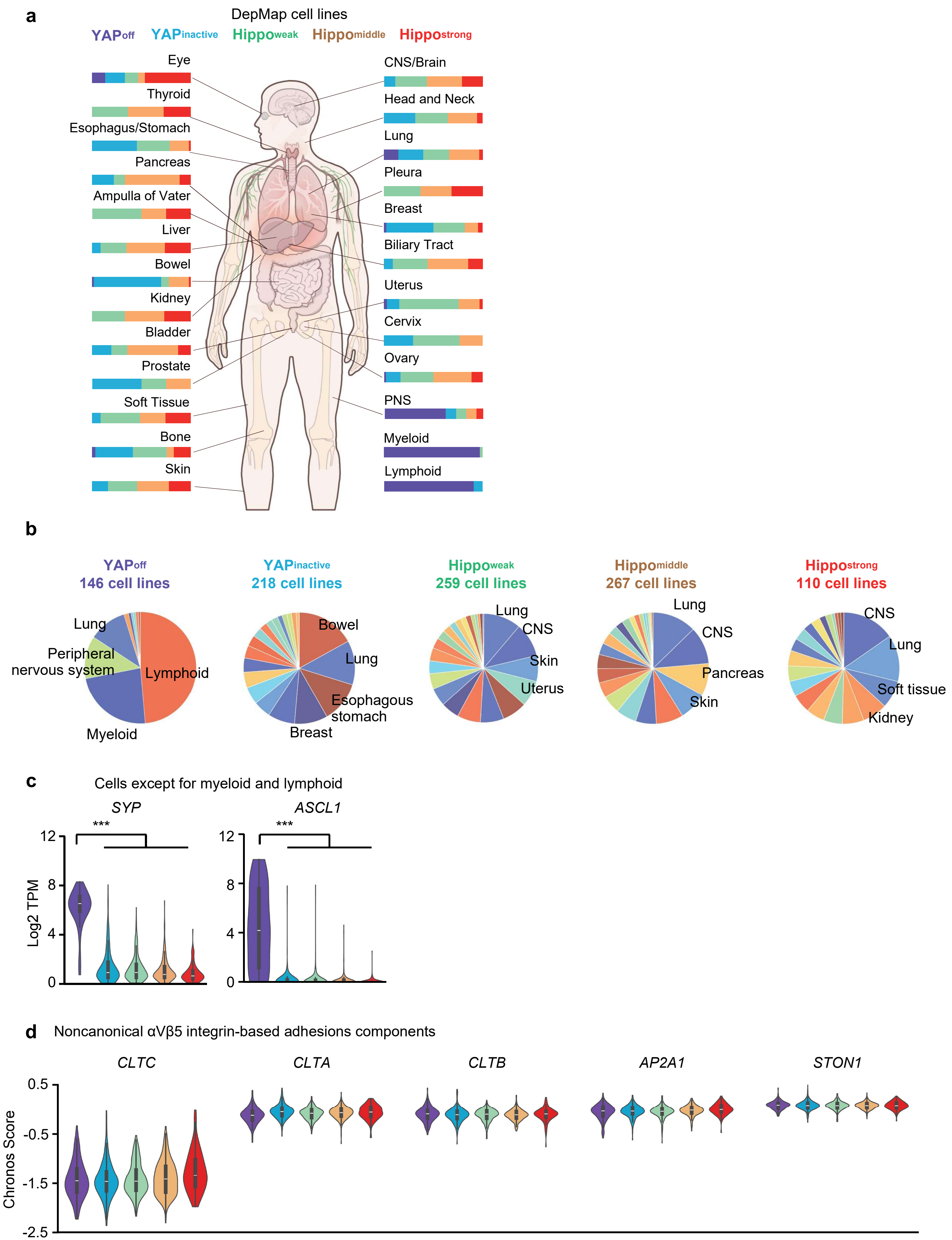

a
DepMap cell lines
YAPoff YAPinactive Hippoweak Hippomiddle Hippostrong
Eye
Thyroid
Esophagus/Stomach
Pancreas
Ampulla of Vater
Liver
Bowel
Kidney
Bladder
Prostate
Soft Tissue
Bone
Skin
CNS/Brain
Head and Neck
Lung
Pleura
Breast
Biliary Tract
Uterus
Cervix
Ovary
PNS
Myeloid
Lymphoid
b
YAPoff
146 cell lines
YAPinactive
218 cell lines
Hippoweak
259 cell lines
Hippomiddle
267 cell lines
Hippostrong
110 cell lines
Lung
Peripheral nervous system
Lymphoid
Myeloid
Bowel
Lung
Esophagous stomach
Breast
Lung
CNS
Skin
Uterus
Lung
CNS
Pancreas
Skin
CNS
Lung
Soft tissue
Kidney
c
Cells except for myeloid and lymphoid
SYP
ASCL1
***
***
Log2 TPM
12
8
4
0
d
Noncanonical αVβ5 integrin-based adhesions components
CLTC
CLTA
CLTB
AP2A1
STON1
Chronos Score
0.5
-0.5
-1.5
-2.5

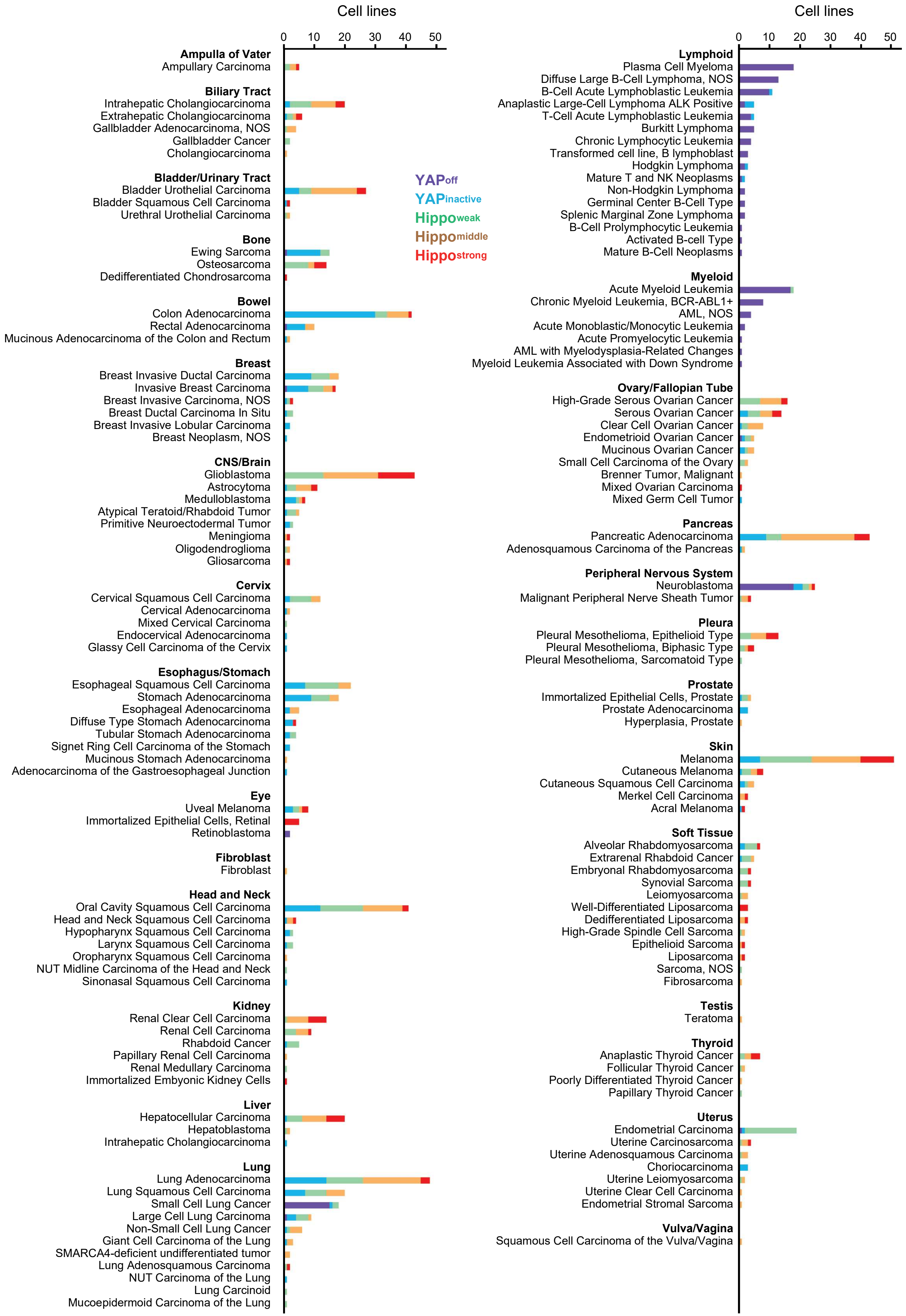

Cell lines
0 10 20 30 40 50
Ampulla of Vater
Ampullary Carcinoma
Biliary Tract
Intrahepatic Cholangiocarcinoma
Extrahepatic Cholangiocarcinoma
Gallbladder Adenocarcinoma, NOS
Gallbladder Cancer
Cholangiocarcinoma
Bladder/Urinary Tract
Bladder Urothelial Carcinoma
Bladder Squamous Cell Carcinoma
Urethral Urothelial Carcinoma
Bone
Ewing Sarcoma
Osteosarcoma
Dedifferentiated Chondrosarcoma
Bowel
Colon Adenocarcinoma
Rectal Adenocarcinoma
Mucinous Adenocarcinoma of the Colon and Rectum
Breast
Breast Invasive Ductal Carcinoma
Invasive Breast Carcinoma
Breast Invasive Carcinoma, NOS
Breast Ductal Carcinoma In Situ
Breast Invasive Lobular Carcinoma
Breast Neoplasm, NOS
CNS/Brain
Glioblastoma
Astrocytoma
Medulloblastoma
Atypical Teratoid/Rhabdoid Tumor
Primitive Neuroectodermal Tumor
Meningioma
Oligodendroglioma
Gliosarcoma
Cervix
Cervical Squamous Cell Carcinoma
Cervical Adenocarcinoma
Mixed Cervical Carcinoma
Endocervical Adenocarcinoma
Glassy Cell Carcinoma of the Cervix
Esophagus/Stomach
Esophageal Squamous Cell Carcinoma
Stomach Adenocarcinoma
Esophageal Adenocarcinoma
Diffuse Type Stomach Adenocarcinoma
Tubular Stomach Adenocarcinoma
Signet Ring Cell Carcinoma of the Stomach
Mucinous Stomach Adenocarcinoma
Adenocarcinoma of the Gastroesophageal Junction
Eye
Uveal Melanoma
Immortalized Epithelial Cells, Retinal
Retinoblastoma
Fibroblast
Fibroblast
Head and Neck
Oral Cavity Squamous Cell Carcinoma
Head and Neck Squamous Cell Carcinoma
Hypopharynx Squamous Cell Carcinoma
Larynx Squamous Cell Carcinoma
Oropharynx Squamous Cell Carcinoma
NUT Midline Carcinoma of the Head and Neck
Sinonasal Squamous Cell Carcinoma
Kidney
Renal Clear Cell Carcinoma
Renal Cell Carcinoma
Rhabdoid Cancer
Papillary Renal Cell Carcinoma
Renal Medullary Carcinoma
Immortalized Embyonic Kidney Cells
Liver
Hepatocellular Carcinoma
Hepatoblastoma
Intrahepatic Cholangiocarcinoma
Lung
Lung Adenocarcinoma
Lung Squamous Cell Carcinoma
Small Cell Lung Cancer
Large Cell Lung Carcinoma
Non-Small Cell Lung Cancer
Giant Cell Carcinoma of the Lung
SMARCA4-deficient undifferentiated tumor
Lung Adenosquamous Carcinoma
NUT Carcinoma of the Lung
Lung Carcinoid
Mucoepidermoid Carcinoma of the Lung
YAPoff
YAPinactive
Hippoweak
Hippomiddle
Hippostrong
Cell lines
0 10 20 30 40 50
Lymphoid
Plasma Cell Myeloma
Diffuse Large B-Cell Lymphoma, NOS
B-Cell Acute Lymphoblastic Leukemia
Anaplastic Large-Cell Lymphoma ALK Positive
T-Cell Acute Lymphoblastic Leukemia
Burkitt Lymphoma
Chronic Lymphocytic Leukemia
Transformed cell line, B lymphoblast
Hodgkin Lymphoma
Mature T and NK Neoplasms
Non-Hodgkin Lymphoma
Germinal Center B-Cell Type
Splenic Marginal Zone Lymphoma
B-Cell Prolymphocytic Leukemia
Activated B-cell Type
Mature B-Cell Neoplasms
Myeloid
Acute Myeloid Leukemia
Chronic Myeloid Leukemia, BCR-ABL1+
AML, NOS
Acute Monoblastic/Monocytic Leukemia
Acute Promyelocytic Leukemia
AML with Myelodysplasia-Related Changes
Myeloid Leukemia Associated with Down Syndrome
Ovary/Fallopian Tube
High-Grade Serous Ovarian Cancer
Serous Ovarian Cancer
Clear Cell Ovarian Cancer
Endometrioid Ovarian Cancer
Mucinous Ovarian Cancer
Small Cell Carcinoma of the Ovary
Brenner Tumor, Malignant
Mixed Ovarian Carcinoma
Mixed Germ Cell Tumor
Pancreas
Pancreatic Adenocarcinoma
Adenosquamous Carcinoma of the Pancreas
Peripheral Nervous System
Neuroblastoma
Malignant Peripheral Nerve Sheath Tumor
Pleura
Pleural Mesothelioma, Epithelioid Type
Pleural Mesothelioma, Biphasic Type
Pleural Mesothelioma, Sarcomatoid Type
Prostate
Immortalized Epithelial Cells, Prostate
Prostate Adenocarcinoma
Hyperplasia, Prostate
Skin
Melanoma
Cutaneous Melanoma
Cutaneous Squamous Cell Carcinoma
Merkel Cell Carcinoma
Acral Melanoma
Soft Tissue
Alveolar Rhabdomyosarcoma
Extrarenal Rhabdoid Cancer
Embryonal Rhabdomyosarcoma
Synovial Sarcoma
Leiomyosarcoma
Well-Differentiated Liposarcoma
Dedifferentiated Liposarcoma
High-Grade Spindle Cell Sarcoma
Epithelioid Sarcoma
Liposarcoma
Sarcoma, NOS
Fibrosarcoma
Testis
Teratoma
Thyroid
Anaplastic Thyroid Cancer
Follicular Thyroid Cancer
Poorly Differentiated Thyroid Cancer
Papillary Thyroid Cancer
Uterus
Endometrial Carcinoma
Uterine Carcinosarcoma
Uterine Adenosquamous Carcinoma
Choriocarcinoma
Uterine Leiomyosarcoma
Uterine Clear Cell Carcinoma
Endometrial Stromal Sarcoma
Vulva/Vagina
Squamous Cell Carcinoma of the Vulva/Vagina

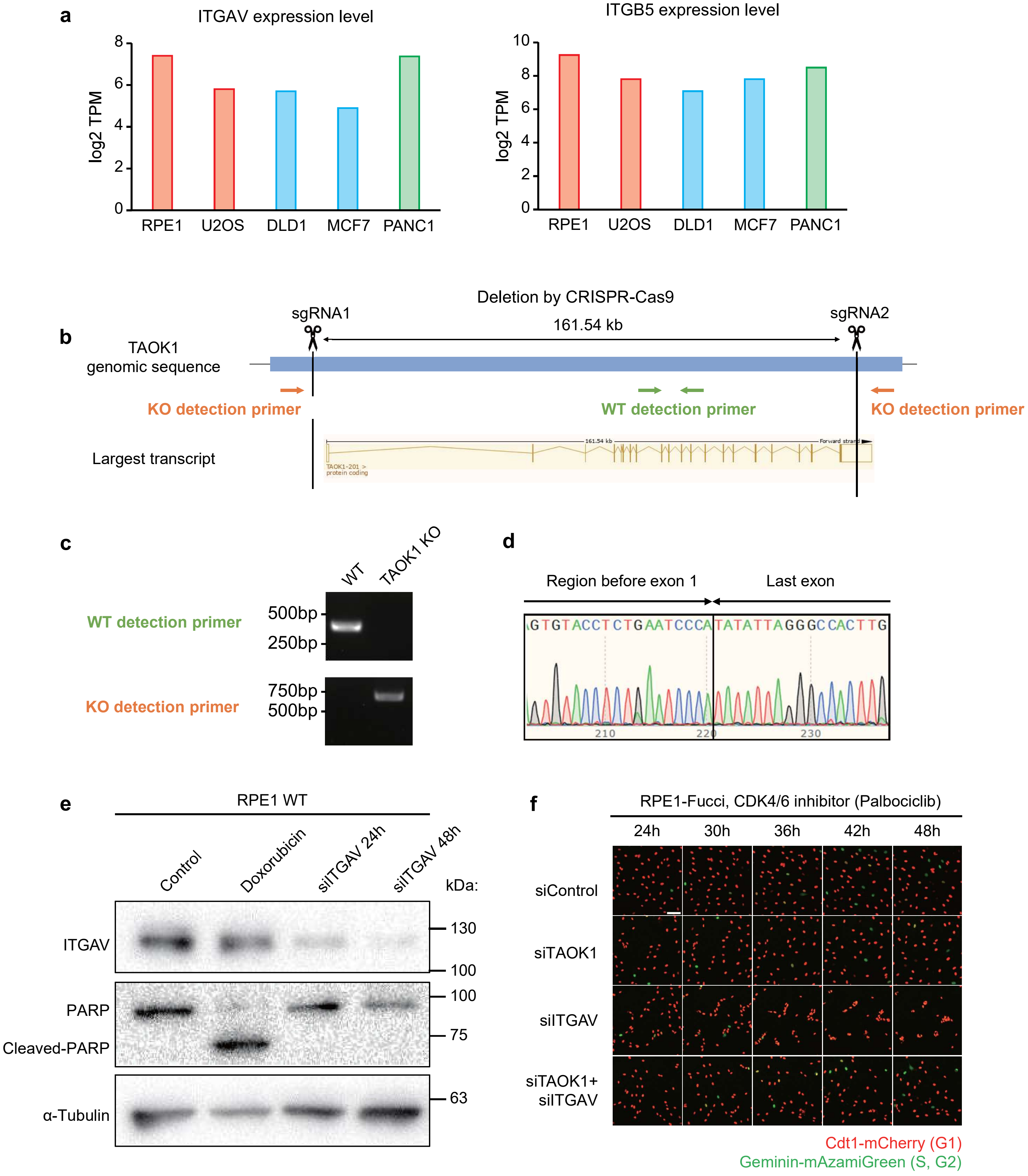

a
ITGAV expression level
ITGB5 expression level
log2 TPM
RPE1
U2OS
DLD1
MCF7
PANC1
b
Deletion by CRISPR-Cas9
sgRNA1
sgRNA2
161.54 kb
TAOK1 genomic sequence
KO detection primer
WT detection primer
KO detection primer
Largest transcript
c
WT
TAOK1 KO
WT detection primer
500bp
250bp
KO detection primer
750bp
500bp
d
Region before exon 1
Last exon
GTGTACCTCTGAATCCCATATATTAGGGCCACTTG
210
220
230
e
RPE1 WT
Control
Doxorubicin
siITGAV 24h
siITGAV 48h
kDa:
ITGAV
130
100
PARP
Cleaved-PARP
100
75
α-Tubulin
63
f
RPE1-Fucci, CDK4/6 inhibitor (Palbociclib)
24h
30h
36h
42h
48h
siControl
siTAOK1
siITGAV
siTAOK1+ siITGAV
Cdt1-mCherry (G1)
Geminin-mAzamiGreen (S, G2)

**a**

**GO enrichment analysis for upregulated genes**

| name | p value |
|---|---|
| MHC class II protein complex | 1.22e-09 |
| ***antigen processing and presentation of exogenous peptide antigen via MHC class II*** | 5.66e-09 |
| antigen processing and presentation of peptide antigen via MHC class II | 1.89e-08 |
| MHC protein complex | 2.10e-08 |
| MHC class II protein complex binding | 7.21e-08 |

**b**

**antigen processing and presentation of exogenous peptide antigen via MHC class II**

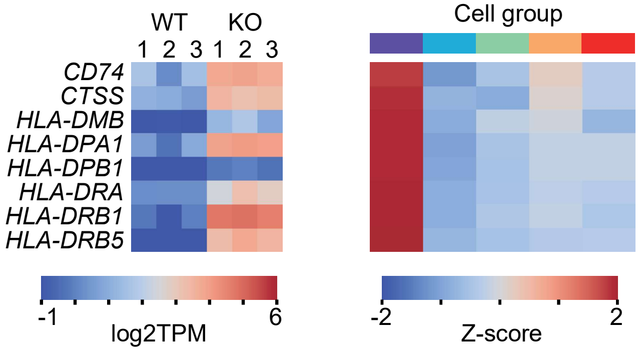

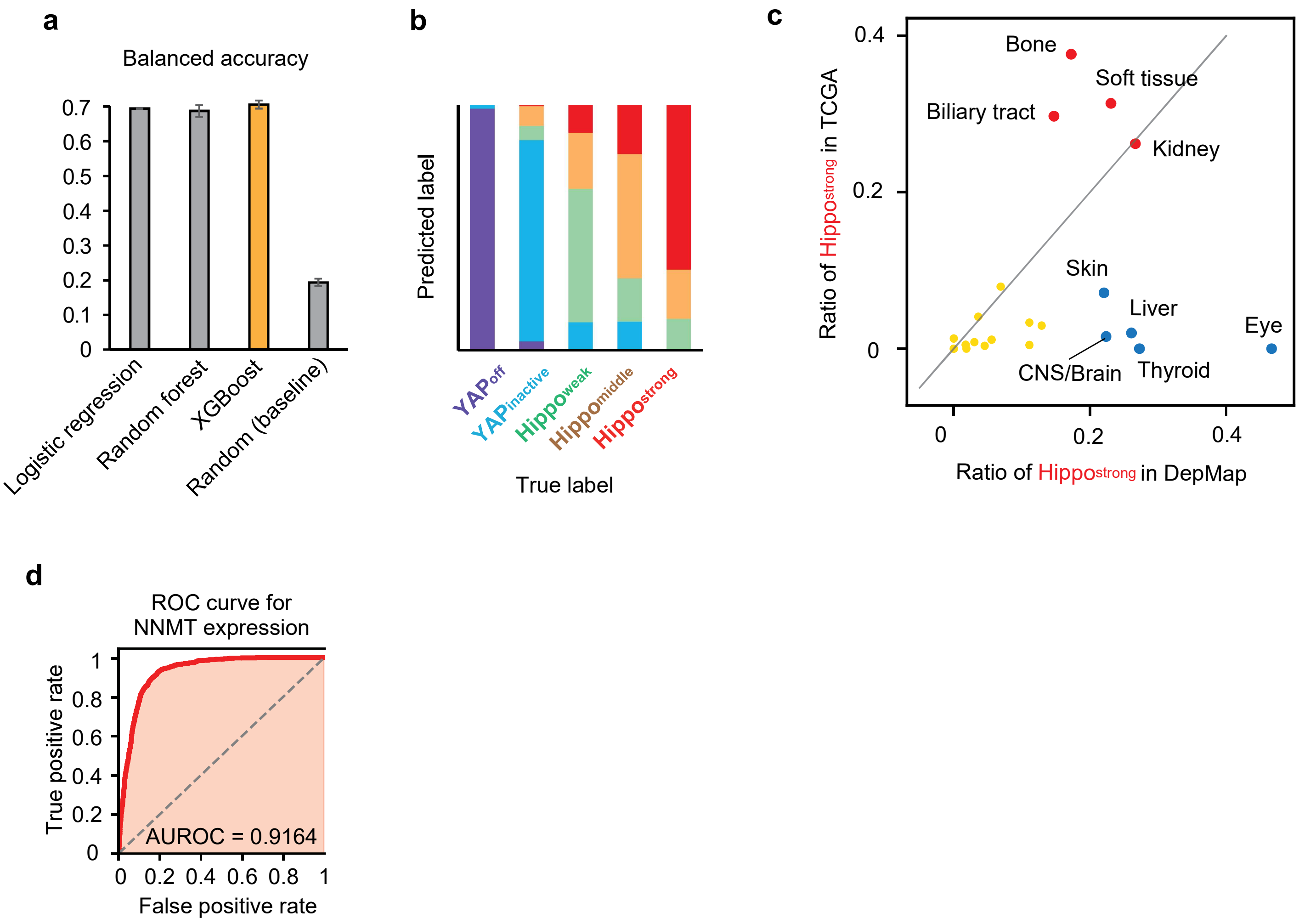

a
Balanced accuracy
0.7
0.6
0.5
0.4
0.3
0.2
0.1
0
Logistic regression
Random forest
XGBoost
Random (baseline)
b
Predicted label
YAPoff
YAPinactive
Hippoweak
Hippomiddle
Hippostrong
True label
c
Ratio of Hippostrong in TCGA
0.4
0.2
0
Bone
Soft tissue
Biliary tract
Kidney
Skin
Liver
Eye
CNS/Brain
Thyroid
0
0.2
0.4
Ratio of Hippostrong in DepMap
d
ROC curve for
NNMT expression
True positive rate
1
0.8
0.6
0.4
0.2
0
AUROC = 0.9164
0
0.2
0.4
0.6
0.8
1
False positive rate
